# Matrix stiffness shifts the endothelial shear stress set point for angiogenic activation

**DOI:** 10.64898/2026.07.22.740002

**Authors:** Laia Gifre-Renom, Ashkan Tabibian, Wolfgang Giese, Femke Bellen, Aernout Luttun, Hans Van Oosterwyck, Elizabeth A.V. Jones

## Abstract

**Aims:** Endothelial cells (ECs) are simultaneously exposed to wall shear stress (SS) from blood flow and substrate stiffness (SFN) from the extracellular matrix, yet how these cues are integrated to shape endothelial behavior remains incompletely understood. We applied an unbiased transcriptomics strategy to define how SS and substrate SFN jointly encode endothelial state transitions and determine angiogenic activation thresholds.

**Methods and Results:** We generated a factorial RNA-Seq dataset of human ECs exposed to 14 combinations of SS (0–40 dynes/cm²) and SFN (1–100 kPa). DESeq2 with likelihood ratio testing identified genes whose expression was significantly associated with SS, SFN, or their interaction. SS was the dominant driver of global transcriptional variation and elicited non-linear transcriptional responses, whereas substrate SFN had a smaller direct effect but significantly modulated the endothelial response to flow. Interaction analyses identified gene programs associated with vascular remodeling, including angiogenesis and migration. Pathway-level analyses revealed that substrate SFN shifts the SS threshold at which angiogenic transcriptional programs become activated, indicating that SFN tunes the endothelial angiogenic set point rather than scaling the response magnitude. Moreover, activated states differed qualitatively across mechanical contexts, reflecting context-dependent reweighting of shared inflammatory, stress, and adaptive/remodeling programs. Finally, siRNA-mediated *YAP1* knockdown confirmed its contribution to SS-SFN-dependent gene regulation.

**Conclusion:** This study provides a systems-level experimental and bioinformatic framework for disentangling multifactorial mechanotransduction in ECs. Although SS predominates in shaping endothelial transcriptomes, substrate SFN critically modulates how ECs interpret flow by shifting the threshold for angiogenic transcriptional activation and reweighting downstream pathways.

**Graphical abstract:** 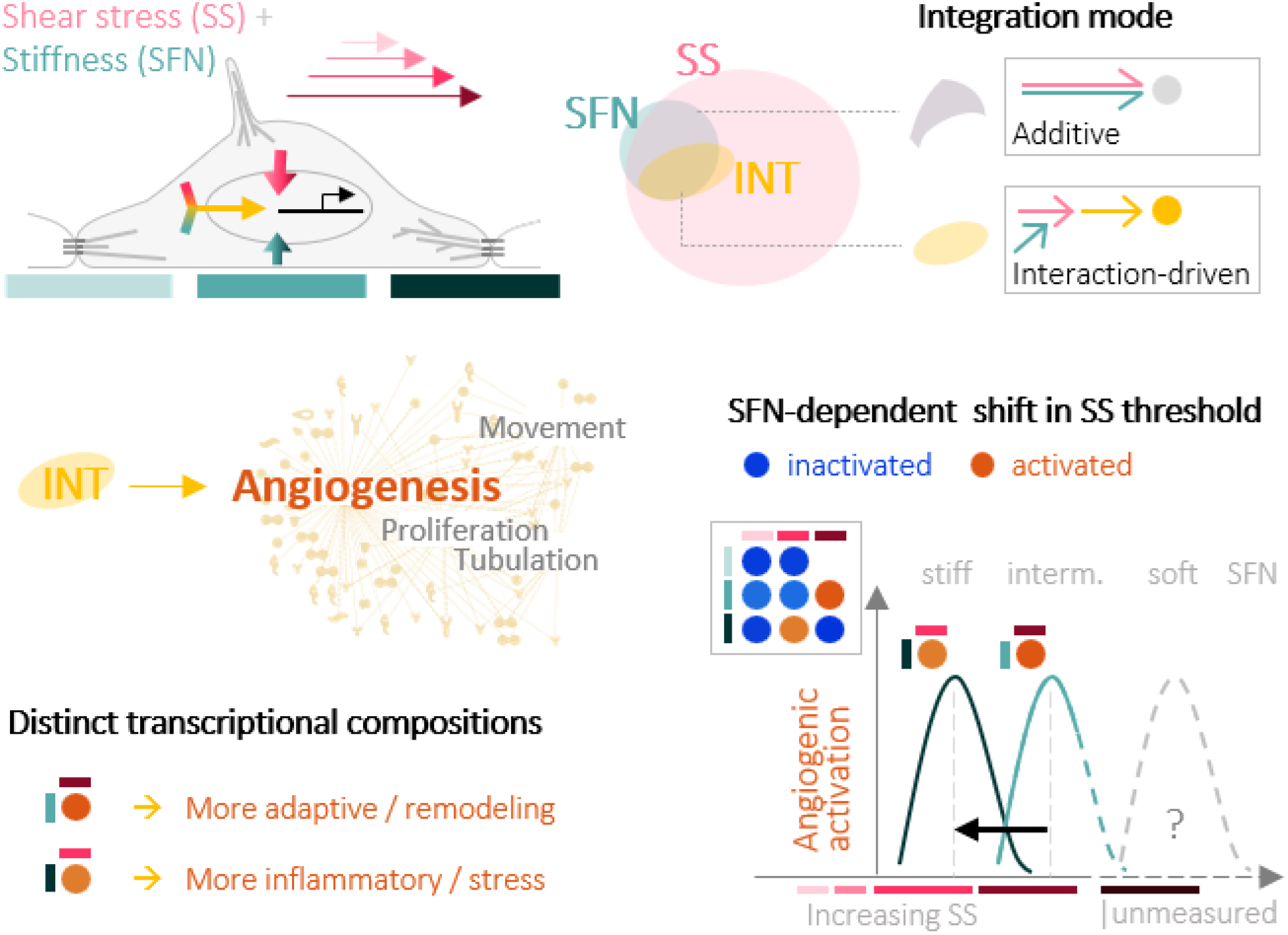

**Translational Perspective:** By systematically combining substrate stiffness and shear stress across physiological and pathological ranges, we provide a reference dataset for vascular mechanobiology. These data enable interpretation of endothelial responses across clinically relevant mechanical environments, including stiffness ranges in different organs (1– brain; 10–muscle or fibrotic liver; 100–aortic valves; kPa), shear stress ranges in different vascular beds (5–veins, 25–capillaries, 40–valves; dynes/cm^2^), and disease-associated changes such as matrix stiffening. Incorporating interactions between mechanical cues may improve the design of *in vitro* vascular models and enhance computational prediction of vascular remodeling.

## 1. Introduction

Endothelial cell (EC) monolayers line the inner surface of the heart, cover cardiac valves, and form the inner lining of blood vessels. As such, they are continuously exposed to mechanical forces generated by blood throughout the cardiovascular system. Mechanotransduction of wall shear stress (SS) is essential for vascular homeostasis, and disturbed flow patterns contribute to atherosclerotic plaque formation, inflammation, and endothelial-to-mesenchymal transition.^1,2^ SS varies considerably throughout the cardiovascular system depending on the type of blood vessel, the organ, and the vessel geometry. Reported values range from as low as 1 dyne/cm^2^ in larger veins^3^ to up to 70 dynes/cm^2^ on the ventricular side of cardiac valves,^4–6^ while estimates in capillaries remain variable, ranging from 5 to 70 dynes/cm^2^.^7^ Importantly, deviations below or above vessel-specific physiological SS ranges can induce pathological endothelial responses. Low SS levels (1-5 dynes/cm^2^) are generally associated with atherosclerosis progression and inflammation,^8,9^ whereas SS levels of 25-40 dynes/cm^2^ are physiological in vascular beds such as aortic valves and arterioles but may become maladaptive outside those vascular environments. As such, a growing body of work suggests that vascular remodeling responses are governed by endothelial SS set points associated with the physiological range of each EC type.^10–12^ For example, human umbilical vein ECs (HUVECs) maintain homeostasis at SS levels between 10-20 dynes/cm^2^ but initiate pro-inflammatory signaling (*e.g.*, NF-κB) when this range is exceeded.^10^ Lymphatic ECs have a lower SS set point (4-6 dynes/cm^2^); however, reducing *VEGFR3* expression by siRNA shifts maximal alignment to 12-15 dynes/cm^2^, similar to that observed in HUVECs.^10^ Therefore, maintaining an appropriate sensitivity to SS is critical for EC function, with implications in vascular disease, including arterial-venous malformations.^13^

In addition to SS, ECs are mechanically coupled to the extracellular matrix (ECM) and surrounding tissue, whose stiffness (SFN) also regulates endothelial physiology. During development, changes in tissue SFN in the cardinal vein induce *GATA2* expression in sprouting ECs, driving their maturation along the lymphatic lineage.^14^ SFN varies widely among organs, ranging from ∼0.5-4 kPa in brain and adipose tissue to ∼10 kPa in lungs, ∼50 kPa in muscle, and up to ∼2 MPa in aortic valves.^15^ In addition, many cardiovascular and metabolic diseases (*e.g.*, hypertrophic cardiomyopathy with heart failure and preserved ejection fraction [HCM-HFpEF] and metabolic dysfunction-associated steatotic liver disease) are associated with tissue stiffening and vascular alterations.^16,17^ Tissue SFN values around 10 kPa have been reported in myocardium from patients with HCM-HFpEF and in fibrotic liver,^16,18^ whereas values around 100 kPa are observed in breast cancer tumors.^19^ Moreover, increased SFN in the fibrotic liver is accompanied by loss of sinusoidal endothelial fenestrations.^20^

Although SS and substrate SFN are sensed simultaneously by ECs, their effects have predominantly been studied in isolation. Only a few studies have examined their combined effects. Kohn *et al.* found that ECs cultured on compliant substrates (2.5 kPa) under 12 dynes/cm^2^ SS exhibit a more atheroprotective phenotype compared with those on stiffer substrates (10 kPa).^21^ Galie *et al.* reached a similar conclusion, finding increased elongation, tight cell-cell junctions and nitric oxide production, and decreased Rho activation at 2.5 kPa under flow.^22^ A transcriptomics study on primary human brain microvascular ECs revealed that inflammatory signatures were activated by increased substrate SFNs, and attenuated when exposed to flow.^23^ However, except for the work by Galie *et al.*, these studies focused on binary conditions, omitting intermediate regimes that are physiologically relevant across vascular beds. Because endothelial responses to mechanical cues are non-linear and governed by set point-like behaviors, a systematic analysis across a broader mechanical landscape is required to capture the gene expression patterns underlying mechanically driven endothelial transition states and determine how substrate SFN reshapes endothelial responses to SS.

Here, we use bulk RNA sequencing in a factorial SS-by-SFN experimental design to systematically investigate endothelial non-linear responses across 14 mechanical conditions, combining different levels of SS (0, 5, 15, 25, 40 dynes/cm^2^) and substrate SFN (1, 10, 100 kPa) for 24 h. Using likelihood ratio testing (LRT) within a negative binomial generalized linear modeling approach combined with functional analyses, we identify interaction-driven transcriptional programs and define how substrate SFN reshapes endothelial transcriptional responses to flow. Our approach reveals non-linear interaction effects, including a SFN-dependent shift in the SS threshold for angiogenic activation that cannot be inferred from single-cue studies. Our findings demonstrate that substrate SFN does not merely scale endothelial responses to flow but instead tunes the SS set point that defines transitions between endothelial functional states.

## 2. Materials and Methods

The complete experimental procedures, including hydrogel fabrication and functionalization, flow system setup (including flow ramp-up), RNA extraction and sequencing, bioinformatic processing, pathway analyses, YAP1 immunostaining and image analysis, endothelial migration assays, *YAP1* knockdown and RT-qPCR protocols, are provided in the Supplementary Methods. The procedures most relevant to the interpretation of the results are summarized below.

### 2.1 Experimental Design

To determine the impact of interactions between SS and substrate SFN on the EC mechanobiology, HUVEC monolayers were cultured on collagen I-functionalized polyacrylamide hydrogels of three substrate SFNs (1, 10, 100 kPa) and exposed for 24h to five levels of steady laminar SS (0, 5, 15, 25, 40 dynes/cm^2^), generating 14 mechanical conditions (the combination of 1 kPa and 40 dynes/cm^2^ was technically unfeasible). RNA sequencing was performed to characterize transcriptional responses to SS, substrate SFN, and their interaction. Transcriptomic findings were complemented with YAP1 nuclear localization analyses, endothelial migration assays, and exploratory *YAP1* knockdown experiments.

### 2.2 Hydrogel preparation at different stiffnesses and EC culture

Polyacrylamide hydrogels with Young’s moduli of 1, 10, or 100 kPa (hydrogel compositions in Table S1) were polymerized on APTS-glutaraldehyde-treated glass slides and functionalized with collagen type I using Sulfo-SANPAH chemistry. HUVECs (C-12208, PromoCell, passage P5-6) were seeded on the hydrogels at 0.25 x 10^6^ cells/hydrogel and incubated for at least 24h before any shear experiment. Detailed hydrogel fabrication, swelling correction (Figure S1), collagen functionalization, and cell culture procedures are provided in the Supplementary Methods.

### 2.3 Shear stress experiments

Hydrogels containing confluent HUVEC monolayers were mounted in custom flow chambers connected to a peristaltic pump and exposed to steady laminar flow for 24h at the indicated SS levels. Static controls were maintained under identical culture conditions without flow. The flow ramp-up protocols (Table S2) and flow system calibration are described in the Supplementary Methods. Cell alignment after flow exposure was confirmed microscopically (Figure S2).

### 2.4 RNA sequencing and differential expression analysis

Total RNA was extracted using QIAzol, and stranded mRNA libraries were sequenced (100 bp paired-end, Illumina NovaSeq 6000). After quality control, reads were trimmed, aligned to the human GRCh38 reference genome, and quantified using standard RNA-Seq pipelines (TrimGalore, HISAT2, featureCounts). Samples with <70% assigned reads were excluded (see Table S3 for replicates, Table S4 for quality control exclusions, and Table S5 for ribosomal genes excluded). Detailed sequencing, preprocessing, alignment, and count-generation procedures are provided in the Supplementary Methods.

Exploratory and differential expression analyses were performed in DESeq2.^24^ Gene counts were fitted to a full negative binomial generalized linear model including batch, SS, SFN, and their interaction. LRTs compared the full model with three reduced models lacking SS, SFN or the interaction term to determine whether inclusion of each term improved model fit (Figure 2A).^25^

When testing the contribution of a main factor (SS or SFN), the interaction term was also removed to preserve model hierarchy, as interactions should not be retained in the absence of their corresponding main effects. The “batch” factor was kept in all models (full and reduced). *P* values were adjusted using the Benjamini-Hochberg procedure, and genes with adjusted *p* < 0.05 were considered significant. Complete LRT results are provided in Supplementary Data 1.

### 2.5 Transcriptomic and functional annotation analyses

Principal component analysis (PCA), Uniform Manifold Approximation and Projection (UMAP), and unsupervised clustering were used to explore global transcriptional structure. Functional interpretation was performed using Ingenuity Pathway Analysis (IPA) and Kyoto Encyclopedia of Genes and Genomes (KEGG). IPA predicts activation or inactivation of biological processes by integrating curated causal relationships with gene expression fold changes obtained from pairwise DESeq2 Wald tests. Because endothelial mechanotransduction studies are traditionally performed under static conditions on rigid plastic or glass, fold changes were calculated relative to the stiffest static condition (100 kPa, static). We extracted fold changes and adj-*p* for all genes and built up a DE data set containing the LRT results for each term and the fold changes and Wald test adj-*p* results across all the condition contrasts for all genes. Complete gene lists for each Wald test result can be found in Supplementary Data 2-3.

KEGG pathway scores were calculated from mean log_2_ fold changes of significantly regulated genes assigned to each pathway. To compare pathway activation between substrate SFN, ΔSFN was calculated as the difference in pathway activation scores between 100 and 10 kPa at each SS level. Positive ΔSFN values indicate greater pathway activation on the stiffer substrate, whereas negative values indicate greater activation on the more compliant substrate. To compare the transcriptional composition of the two angiogenesis-associated states (25 dynes/cm² at 100 kPa and 40 dynes/cm² at 10 kPa), ΔMech was calculated as the difference in gene log₂ fold change between both conditions. Positive ΔMech values indicate higher expression at 40 dynes/cm²/10 kPa, whereas negative values indicate higher expression at 25 dynes/cm²/100 kPa.

KEGG pathways ranked by ΔSFN and genes contributing to the divergence between angiogenic profiles ranked by ΔMech are provided in Supplementary Data 4. Additional computational details on PCA, UMAP, clustering, IPA, KEGG, and pathway scoring are described in the Supplementary Methods.

### 2.6 Endothelial migration assay

Migration was quantified on HUVECs seeded on fibronectin-coated hydrogels of SFNs of 0.5, 3.2, 4.5, 10, and 35 kPa. Fluorescently labeled HUVECs were tracked during 5h of exposure to 1 dyne/cm^2^ SS, using live confocal microscopy, and migration rates were calculated from individual cell trajectories using tracking algorithm of Imaris 9.1 (Bitplane) as previously described.^26^ Detailed hydrogel preparation (Table S6), live-cell imaging, and tracking procedures are described in the Supplementary Methods.

### 2.7 Active YAP1 localization

Active YAP1 nuclear localization was quantified by immunofluorescence across all SS-SFN conditions using confocal microscopy. Relative nuclear localization was expressed as the ratio of nuclear to whole-cell fluorescence intensity. Antibody information, staining protocol, image acquisition, and quantification procedures are provided in the Supplementary Methods.

### 2.8 YAP1 knockdown

To explore the contribution of YAP1 to interaction-responsive transcription, HUVECs were transfected with *YAP1*-targeting or non-targeting siRNA and exposed to representative mechanical conditions. Expression of selected candidate genes (Supplementary Data 5) was quantified by RT-qPCR. Electroporation protocol, culture conditions, candidate gene selection criteria, and qPCR details are provided in the Supplementary Methods, Table S7 (primer list), and Supplementary Data 5 (ΔCt, ΔΔCt, and confidence interval values).

### 2.9 Statistical Analysis

Statistical analyses were performed in R (v. R-4.3.1). RNA-Seq analysis was conducted on 47 samples (*n* = 3 to 5 samples per condition; *n* = 2 for 1 kPa at 15 dynes/cm^2^) using the LRT and Wald test methods in DESeq2 as previously described. P-values were adjusted for multiple testing using the Benjamini-Hochberg method, and adjusted *p*-values < 0.05 were considered statistically significant. Quadratic and linear regression curves on gene count plots were obtained using the geom_smooth function in ggplot2 in R.

Gene expression functional analyses were run in IPA (v. 107193442) and KEGG. Prediction statistical details for IPA are available online. KEGG pathway analyses were based on pathway-level aggregation of log_2_ fold changes from significant pathway-member genes, using adjusted *p* < 0.05 and a minimum pathway size of 3 genes.

YAP1 localization was analyzed by two-way ANOVA followed by Tukey’s honestly significant difference (HSD) post hoc test. For qPCR experiments, statistical analyses were performed on ΔCt averaged at the biological replicate level. *YAP1* knockdown effects were quantified as differences between si*YAP1* and NT control within each condition. Because the experiment was exploratory and sample sizes were small (*n* = 3–4 biological replicates per condition), effect sizes were summarized using bootstrap resampling to estimate 95% confidence intervals.

## Data availability

The RNA-Seq raw data, as well as LRT and Wald test results can be found in the NCBI Gene Expression Omnibus (GEO) repository with accession number GSE262429 (available after publication in journal). All other data are present in the paper and/or the Supplementary Materials.

## 3. Results

### 3.1 Shear stress dominates endothelial transcription variation

To investigate the independent and combined effects of SS and substrate SFN on endothelial transcription, we performed bulk RNA sequencing on HUVECs cultured under a factorial design comprising five SS levels (static, 5, 15, 25, and 40 dynes/cm^2^) and three substrate SFNs (Young’s modulus of 1, 10, and 100 kPa; Figure 1A). The 40 dynes/cm^2^ SS at 1 kPa SFN condition was technically unfeasible and excluded. After quality control, 47 samples were retained for analysis.

**Figure 1:**
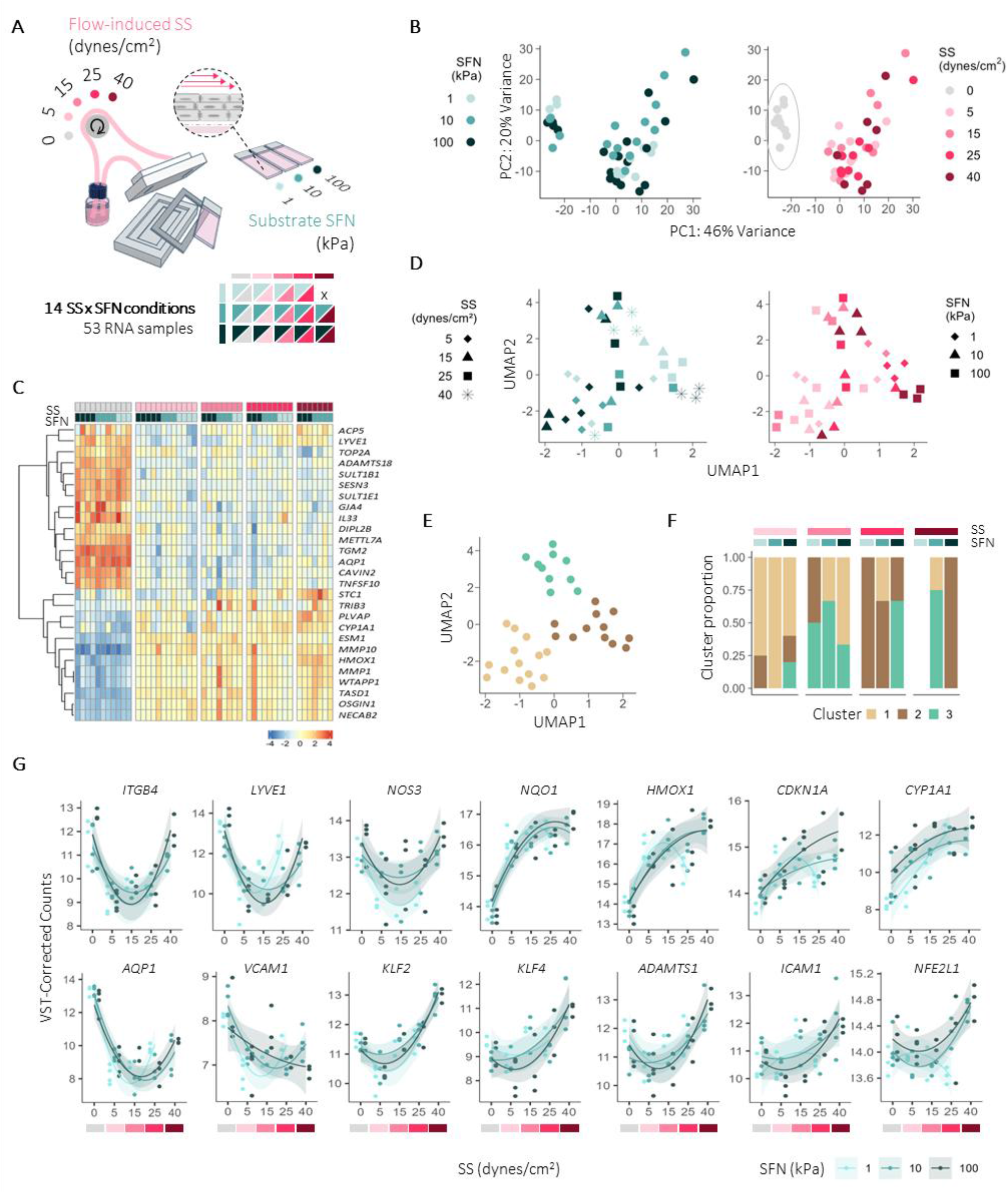
Transcriptomic overview of the endothelial cells exposed to 14 combinations of shear stress (SS) and substrate stiffness (SFN). A, Experimental design and parallel-plate flow chamber. Pink palette, SS (gray for static); turquoise palette, SFN. B, Principal component analysis (PCA) of all samples colored by SFN (top) and by SS (bottom) Static samples are outlined. C, Heatmap of some of the genes with highest variation across conditions (Variance Stabilizing Transformation [VST]-normalized counts centered per gene across samples). Columns indicate individual samples. D, Uniform Manifold Approximation and Projection (UMAP) of flow-exposed samples after exclusion of static SS samples, colored by SFN and with shapes indicating SS (left) and *vice versa* (right). E, UMAP colored by unsupervised k-means clusters (k = 3). F, Cluster composition across flow conditions. G, Gene count dynamics (VST-corrected) of representative flow-responsive genes across SS levels, colored by SFN. Curves represent quadratic regression fits (“lm” smoothing method with formula y ∼ poly(x, 2)) with shaded standard errors. *n* = 3-5 samples/condition (n = 2 for 1 kPa at 15 dynes/cm^2^).

PCA identified SS as the main source of transcriptional variation, with static samples separating from all flow conditions along PC1 (46% variance; Figure 1B), irrespective of SFN. The same separation was evident in the heatmap of the 50 most variable genes (Figure 1C), including established flow-responsive genes^27^ involved in oxidative stress, endothelial identity, and ECM remodeling (e.g., *CYP1A1*, *HMOX1*, *MMP1*/*MMP10, AQP1*, and *LYVE1*).

To resolve transcriptomic variation among flow conditions beyond the dominant separation of static samples, we repeated the analysis after excluding static samples and performed UMAP (Figure 1D). Low SS (5 dynes/cm^2^) formed a distinct cluster that matched unsupervised cluster 1 (Figure 1F), whereas higher SS conditions scattered across the UMAP, dominating clusters 2 and 3 (Figure 1F), indicating greater transcriptomic heterogeneity. Partial grouping of specific SS-SFN combinations (*e.g.*, 40 dynes/cm^2^ at 10 kPa, 25 dynes/cm^2^ at 100 kPa and 15 dynes/cm^2^ at 10 kPa, vs. 25 dynes/cm^2^ at 1 kPa and 40 dynes/cm^2^ at 100 kPa) suggested that both SS and SFN jointly shape transcriptomic organization without either factor alone producing complete segregation (Fig 1E-F).

We next examined individual SS-responsive genes to verify known flow-dependent signatures in our dataset (Figure 1G). By virtue of the factorial design, we examined gene expression patterns across SS and SFN conditions, identifying mostly non-linear responses across increasing SS levels, revealing transcriptome-wide patterns not previously characterized. Here, we identified (i) symmetrical U-shaped responses, with similar expression under static and high SS conditions but different at intermediate SS levels (*e.g.*, *ITGB4*, *LYVE1*, and *NOS3*, upregulated under static and high SS levels); (ii) asymmetrical responses showing the largest difference between static and flow conditions (*e.g.*, *NQO1*, *HMOX1*, *CDKN1A*, CYP1A1, upregulated with SS; *AQP1* and VCAM1, downregulated); and (iii) shear-regime-dependent responses with distinct expression at low SS (5 dynes/cm^2^) relative to both static and, most prominently, the highest SS level (*e.g., KLF2*, *KLF4*, *ADAMTS1*, *ICAM1*, and *NFE2L1*, downregulated at low SS and upregulated with increasing SS). Consistent with the PCA, SFN effects were less pronounced, and SFN-dependent patterns could not be resolved at this stage of the analysis (Figure S3). Nevertheless, known SFN-responsive genes (*e.g.*, *PXN*, *RHOF*, *RHOT2*, *MAP2K3*, and *CDH5*)^28–30^ increased with SFN (Figure S4). Also upregulated with SFN was *LIMS2*, which encodes a focal adhesion protein that interacts with integrin-linked kinase (ILK), known to be modulated by changes in SFN.^31^ In contrast, several docking protein genes like *DOCK1*, *DOCK4* and *DOCK10*, and *SEPTIN10* decreased with increasing SFN. Interestingly, many SFN-responsive genes were associated with cell migration and/or cytoskeletal remodeling.^28,32–34^

### 3.2 The SS-SFN interaction significantly modulates genes in ECs

To assess the contribution of SS, SFN, and their interaction to endothelial gene expression, we fitted the expression data using a full DESeq2 model including SS, SFN, and their interaction (INT; SS x SFN),^24,35^ and performed LRTs by comparing the full model with three reduced models lacking each term (Figure 2A).^25^ Unlike a Wald test, which compares two specific conditions, the LRT tests whether an entire factor is needed to explain the expression patterns across all conditions. Genes significant in each LRT were therefore considered associated with the corresponding model term, yielding 9,486 SS-associated genes, 1,258 SFN-associated genes, and 603 interaction-associated genes (Figure 2A).

**Figure 2:**
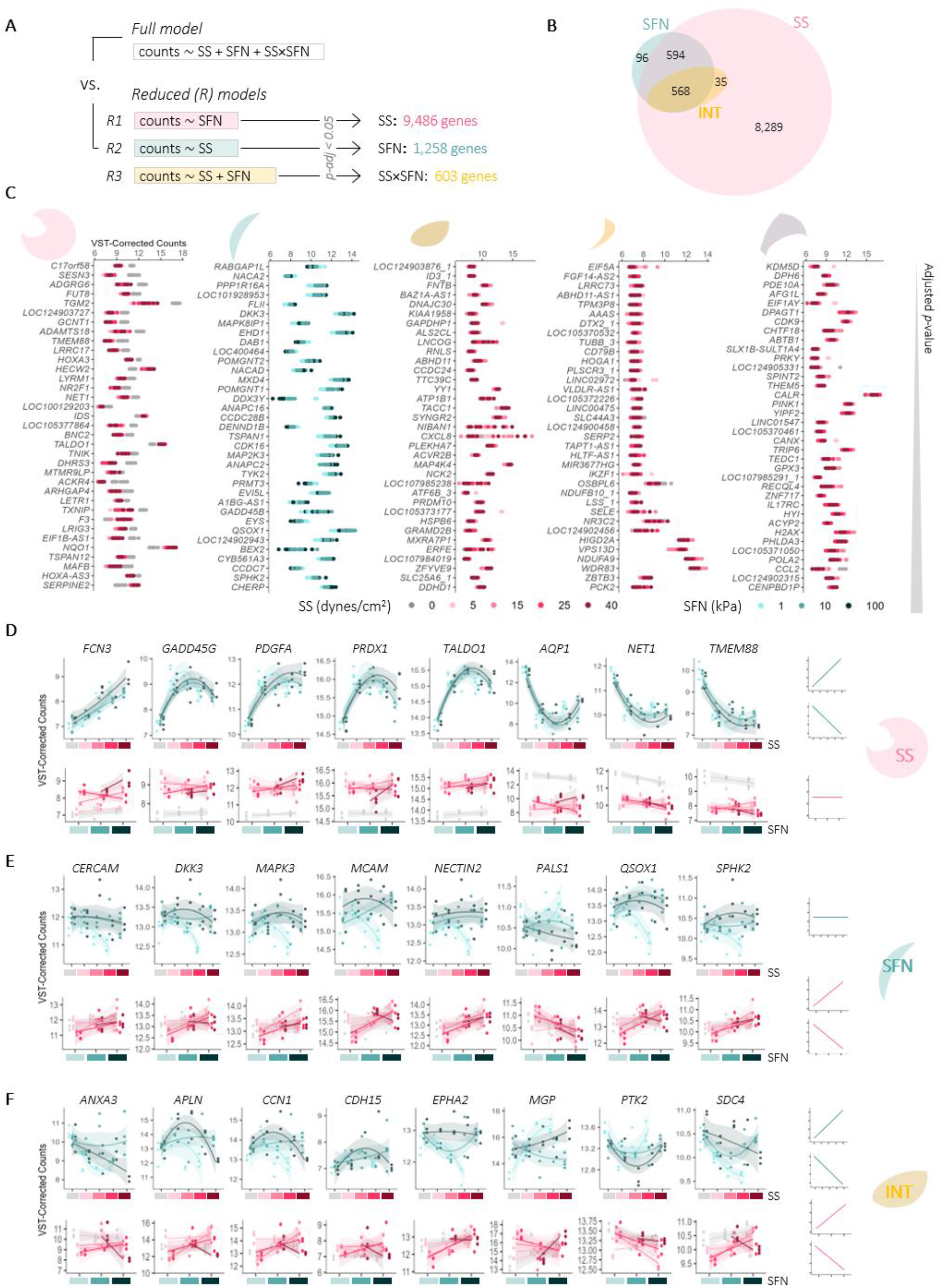
Identification of shear stress (SS)-, substrate stiffness (SFN)-, and interaction (INT, SS×SFN)-responsive genes. **A**, Likelihood ratio testing strategy comparing the full DESeq2 model with reduced models for SS, SFN, and interaction effects. Numbers indicate significant genes (adj-*p* < 0.05).B, Venn diagram showing the overlap between the 3 significant gene lists. C, Top-ranked genes from each Venn diagram subset as VST-corrected expression profiles. Complete gene lists are provided in Supplementary Data 1. D-F, Representative expression patterns for SS-unique (D), SFN-unique (E), and INT-unique (F) responsive genes. Top panels show expression across SS (colored by SFN); bottom panels show expression across SFN (colored by SS). Curves represent fitted regressions (quadratic for SFN across SS; linear for SS across SFN) with shaded standard errors. *n* = 3-5 samples per condition (*n* = 2 for 1 kPa at 15 dynes/cm²). Simplified schematics of pure SS, SFN, and interaction effects, shown as linear patterns for clarity, accompany each panel.

Most SS-associated genes were unique to SS (8,289; SS-unique), whereas only 96 genes were unique to SFN (SFN-unique; Figure 2B). Among genes associated with both cues, 568 also showed significant SSxSFN interactions (overlap-INT), while 594 did not (overlap-noINT), indicating additive rather than interactive effects. Conversely, 35 interaction-associated genes were absent from the SFN list (INT-35), suggesting that SFN influenced their expression only in combination with SS.

Representative genes from each subset are shown in Figure 2C. SS-unique genes included regulators of antioxidant processes (*SESN3*, *TALDO1*, *NQO1*), ECM remodeling (*TGM2*, *ADAMTS18*), cell adhesion (*CLDN10*, *CLDN11*), and transcription (*NR2F1*) (Figure 2C, first plot). SFN-unique genes included mediators of cytoskeleton organization (*FLII*, *PPP1R16A*), vesicle transport (*RABGAP1L*, *CDK16*), and kinase signaling (*MAP2K3*, *TYK2*, *GADD45B*) (Figure 2C, second plot). Top genes shared by SS and SFN with a significant interaction (overlap-INT) were associated with cytoskeletal dynamics (*ATP1B1*, *TAL1*, *SYNGR2*, *PICK1*, *TRIM46*), inflammation (*CXCL8*, *IL6*, *MAP4K4*, *PTX3*, *TRAFD1*), and enzymes involved in hypertension and hydrolysis (*RNLS*, *ABHD11*) (Figure 2C, third plot). Among the INT-35 genes, around 30% were related to mitochondrial respiration (*HOGA1*, *PLSCR3_1*, *NDUFB10*, *HIGD2A*, *VPS13D*, *NDUFA9*, *PCK2*), as well as to cholesterol metabolism (*LSS_1*, *OSBPL6*), cytoskeleton dynamics (*EIF5A*, *TUBB_3*), transmembrane transport (*SLC44A3*), immune responses (*CD79B*, *SELE*), and transcription (*IKZF1*, *ZBTB3*) (Figure 2C, fourth plot); however, these genes generally showed lower read counts than the other subsets. Overlap-noINT genes included regulators of protein folding (*CALR*, *CANX*), mitochondrial function (*AFG1L*, *THEM5*, *PINK1*), transcription (*CDK9*), antioxidant enzymes (*GPX3*), and chemotaxis (*CCL2*) (Fig 2C, fifth plot), suggesting that SS-SFN additive effects may reinforce core endothelial homeostatic functions. Supporting literature of the mentioned gene examples is provided in Table S8.

As expected, expression profiles reflected the LRT classification (Figure 2D-F): SS-unique genes varied predominantly across SS (Figure 2D, top), SFN-unique genes varied across SFN (Figure 2E, bottom), and interaction-associated genes varied across both mechanical cues (Figure 2F).

### 3.3 Substrate SFN shifts the SS threshold at which ECs activate angiogenesis transcriptional programs

To assess global transcriptional changes, we compared gene expression at each SS level with its corresponding static condition (Figure 3A) or with the stiffest static condition (100 kPa), which approximates the glass-like substrate commonly used in the field (Figure 3B). In both analyses, the total number of DEGs peaked at 5 dynes/cm^2^ SS on 1 kPa substrates but shifted to 15 dynes/cm^2^ on 10 and 100 kPa substrates. Notably, however, the pattern of DEGs differed significantly based on SFN, regardless of which condition was used as control.

**Figure 3:**
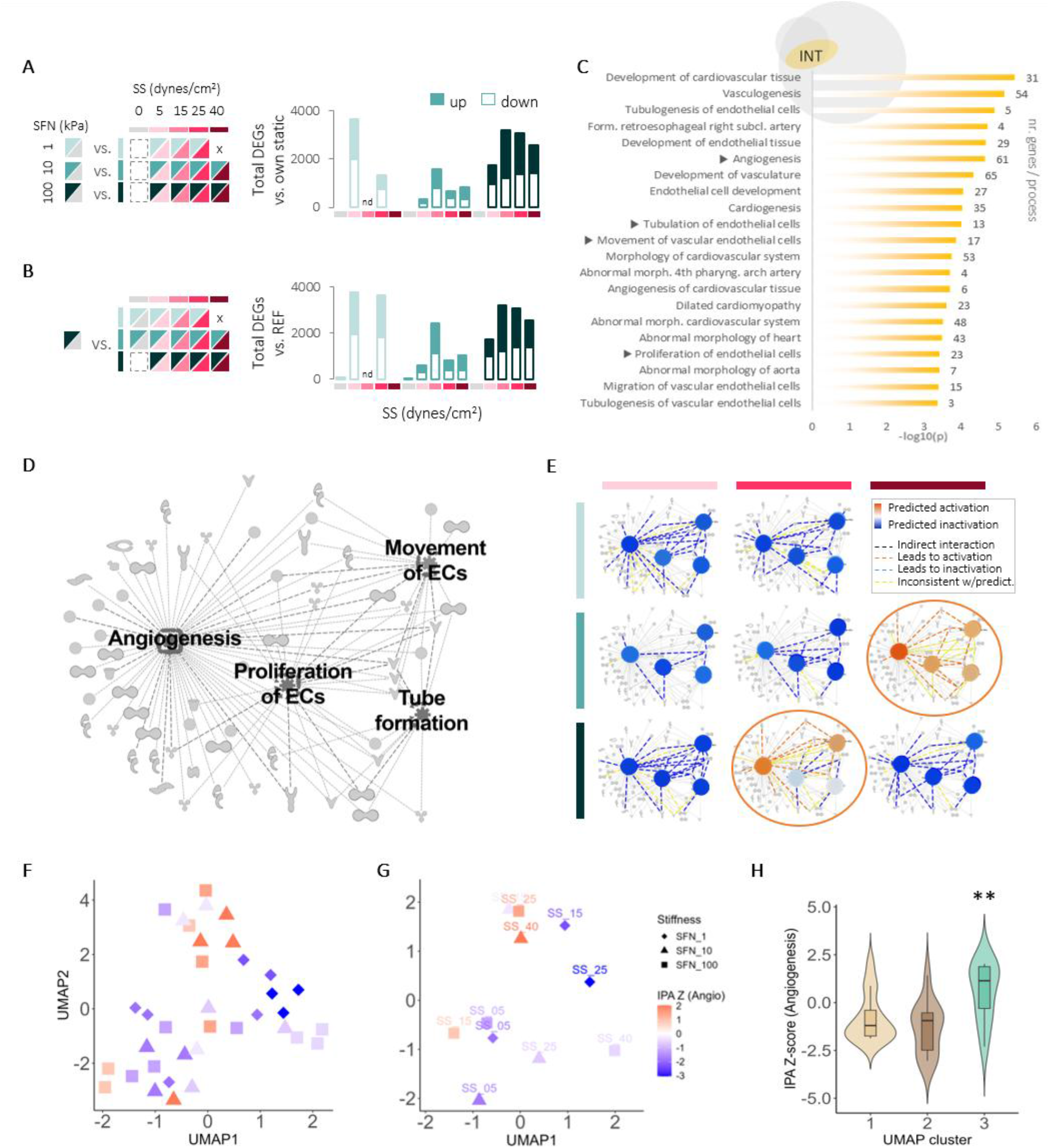
Predicted modulation of cardiovascular processes by interaction DEGs. A,B, Total differentially expressed genes (DEGs) across conditions relative to (A) corresponding static condition within each SFN or (B) the stiffest static condition (100 kPa) (Wald tests, adj-*p* < 0.05). C, Top “cardiovascular system development and disease” processes identified by IPA from the interaction-associated genes. Processes are ordered by -log(*p*); numbers indicate supporting genes. Arrowheads denote the four selected processes. D, Summarized network of the selected processes. Gene names and shape codes are provided in Figure S5. E, Predicted activation states of the selected processes across conditions (IPA Z-scores). Activation: Z > 2 (orange); inactivation: Z < -2 (blue). Predictions for 15 dynes/cm^2^ SS were omitted because the 1 kPa condition included only two samples; the complete overview is shown in Figure S6 and Supplementary Movie 1. F, UMAP of flow-exposed samples colored by angiogenesis activation Z-score. G, UMAP centroids colored by angiogenesis activation Z-score. H, Violin plots with overlaid boxplots showing angiogenesis Z-score distributions across the three UMAP-derived k-means clusters (one-way ANOVA, 36 samples, *p* = 0.0029). *n* = 3-5 samples per condition (*n* = 2 for 1 kPa at 15 dynes/cm^2^).

IPA CORE analysis was performed on the 603 interaction-associated genes to predict regulated biological processes based on curated gene-function relationships. Unlike enrichment analyses, IPA predicts process activation by integrating gene fold changes with curated information on whether each gene activates or inhibits a given process. The top molecular and cellular functions obtained by IPA were “cell death and survival”, and “cellular movement”, followed by “cellular response to therapeutics” and “carbohydrate metabolism”. The top physiological categories were “embryonic development”, “cardiovascular system development and function”, and “tissue development” (Table S9).

From the top 20 cardiovascular-related processes (Figure 3C), we focused on endothelial and cardiovascular functions, excluding embryonic-development terms likely reflecting the umbilical origin of HUVECs. Where redundant processes shared many supporting genes (*e.g.,* tubulation and tubulogenesis of ECs), we retained that supported by the largest gene set. Angiogenesis, tubulation of ECs, movement of vascular ECs, and proliferation of ECs were selected for further analysis (Figure 3C). A network of the selected processes and their supporting genes was generated (Figure 3D; full network in Figure S5) and overlaid with gene fold changes relative to the stiffest static condition to predict process activation across conditions.

The comparison of the resulting Z-score predictions across all conditions revealed distinct activation patterns (Figure 3E; full network-per-condition panel in Figure S6). All three SFNs showed the four processes generally inactivated (blue nodes) under low SS levels (5 and 15 dynes/cm^2^). Notably, increasing SS promoted a shift toward activation (orange nodes), with the transition point depending on substrate SFNs. At 1 kPa, none of the processes became activated at any SS level. At 10 kPa, activation occurred only at 40 dynes/cm^2^. On 100 kPa substrates, processes showed a trend toward activation at 25 dynes/cm^2^ SS before reverting to inactivation at the highest SS level.

Related processes showed similar behavior. Cellular homeostasis, which includes genes involved in stress responses and survival mechanisms, shifted from activation at 40 dynes/cm^2^ SS on 10 kPa to 15 and 25 dynes/cm^2^ SS on 100 kPa substrates (Figure S7), while microtubule dynamics mirrored EC movement (Figure S7). Together, these findings suggest that increasing substrate SFN can sensitize ECs to SS, lowering the SS threshold required for process activation, rather than simply modulating the magnitude of response. To determine whether angiogenic activation contributed to the heterogenic UMAP clustering at high SS, we overlaid the Z-scores for the cardiovascular process showing the strongest response, angiogenesis, onto the UMAP (Figure 3F, G). One of the two k-means clusters corresponded predominantly to an active angiogenic transcriptional state (cluster 3, Figure 3H and Figure 1E), indicating that the activation of angiogenesis was a major driver of transcriptional divergence under elevated SS.

### 3.4 Distinct SS-SFN combinations shape pathway balances underlying angiogenesis-associated transcriptional states

To validate the IPA findings using an independent annotation framework, we performed KEGG-based pathway analysis on the two conditions transcriptionally predicted to activate angiogenesis: 25 dynes/cm^2^ SS at 100 kPa, and 40 dynes/cm^2^ SS at 10 kPa substrates. For each SS level, we compared pathway activation (mean log_2_FC of significantly regulated genes assigned to a given pathway) between 10 and 100 kPa substrates (ΔSFN; Figure 4A-B), allowing us to determine whether a pathway was preferentially activated on one SFN or the other at a given SS. At 25 dynes/cm^2^ SS (Figure 4A-B, left), most pathways showed preferential activation on 100 kPa than on 10 kPa (positive ΔSFN; pathways lying above the identity line that indicates ΔSFN=0). These included inflammatory and stress-associated pathways (NF-κB, TNF, cytokine-cytokine receptor interaction, apoptosis and ferroptosis), endothelial activation and adaptation signaling pathways (MAPK, PI3K-Akt, and mTOR), and adhesion- and trafficking-related pathways (cell adhesion molecules, endocytosis, and phagosome). Only a few pathways, including Notch and adherens junction signaling, were relatively more active on 10 kPa. At 40 dynes/cm^2^ SS (Figure 4A-B, right), this pattern was reversed. Most pathways were more activated on the compliant substrate (10 kPa), including ligand-receptor signaling, inflammatory signaling (NF-κB, TNF, cytokine signaling), cell cycle, cellular senescence, cGMP-PKG, motor proteins, and adaptation and metabolism pathways (AMPK, HIF-1, and FoxO signaling). Conversely, ECM-receptor interaction and TGF-β signaling displayed a relatively higher activation on 100 kPa.

**Figure 4:**
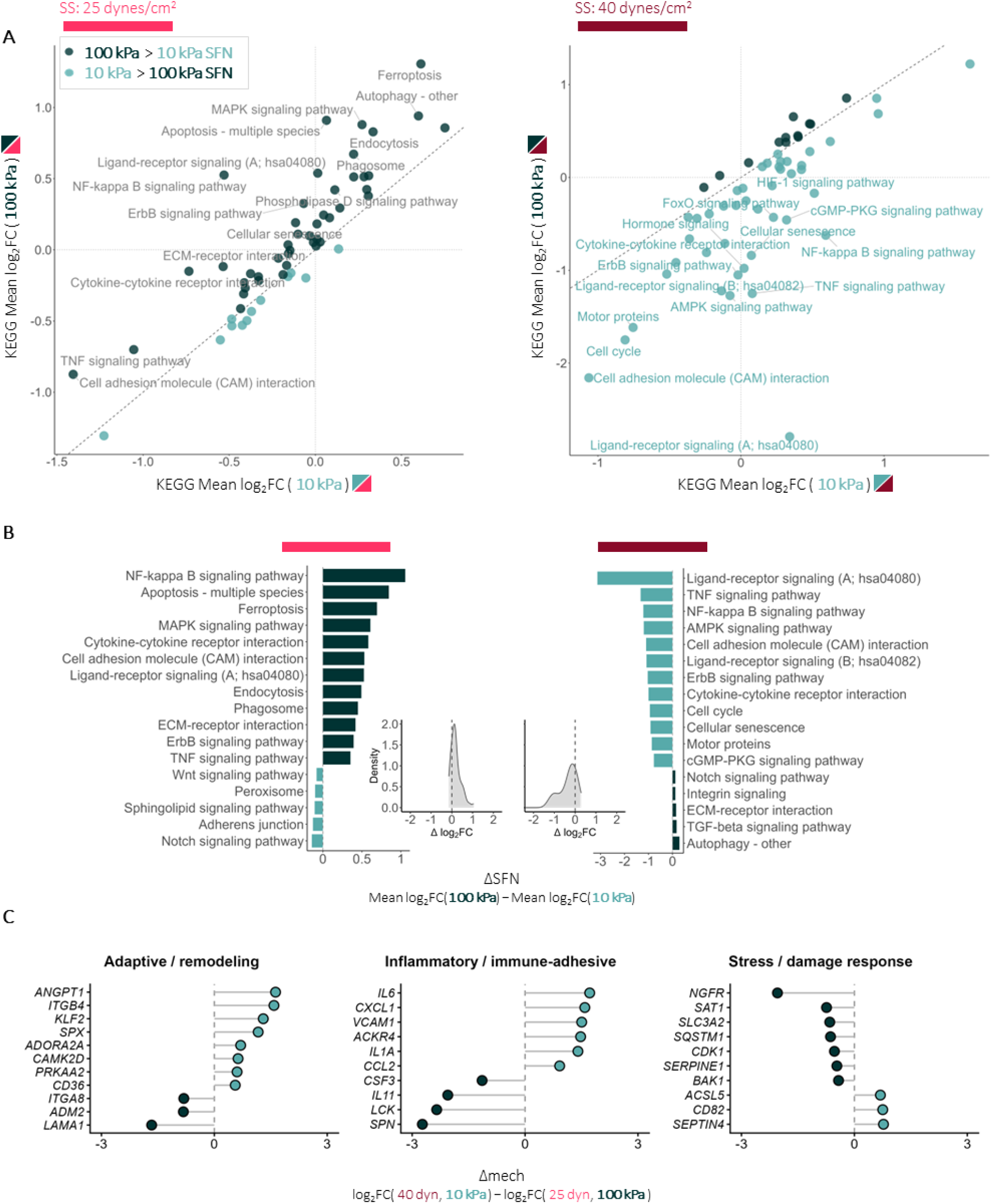
Distinct transcriptional programs underlie angiogenic activation states under different SS-SFN combinations. A, Scatter plots comparing KEGG pathway activation scores (mean log_2_FC of significantly regulated genes assigned to each pathway) between 100 and 10 kPa SFN at 25 dynes/cm^2^ (left) and 40 dynes/cm^2^ SS (right). Each dot represents one KEGG pathway. Dashed identity lines indicate equal pathway activation between substrate SFNs; pathways above the line, preferentially activated at 100 kPa (dark green); pathways below the line, preferentially activated at 10 kPa (light green). Pearson correlation coefficients: *r* = 0.87 (*p* = 4.8×10^-18^) at 25 dynes/cm^2^ and *r* = 0.71( *p* = 9.6×10^-10^) at 40 dynes/cm^2^. B, Most differentially activated KEGG pathways between 100 and 10 kPa at 25 dynes/cm^2^ (left) and 40 dynes/cm^2^ SS (right), expressed as ΔSFN = mean log_2_FC (100 kPa) – mean log_2_FC (10 kPa). Positive values indicate greater activation at 100 kPa and negative values greater activation at 10 kPa. Density plots show distribution of ΔSFN values, with the dashed line indicating ΔSFN = 0. The complete pathway list is provided in Supplementary Data 4. C, Genes contributing most strongly to the transcriptional divergence between the two angiogenic states (25 dynes/cm^2^ at 100 kPa and 40 dynes/cm^2^ at 10 kPa). Genes were grouped into three functional categories as described in Methods and detailed in Supplementary Data 4.

These findings support the IPA prediction that substrate SFN shifts the SS threshold for endothelial transcriptional activation. However, they also indicate that the two angiogenesis-associated states were not transcriptionally identical.

To identify the molecular basis of these differences, we directly compared the genes associated with the preferentially activated pathways between the two angiogenesis-associated conditions (25 dynes/cm² at 100 kPa and 40 dynes/cm² at 10 kPa; Figure 4C). Genes were ranked by the difference in their log_2_FC between conditions (ΔMech), with positive values indicating higher expression at 40 dynes/cm^2^ at 10 kPa and negative values indicating higher expression at 25 dynes/cm^2^ at 100 kPa. To facilitate interpretation, pathway-associated genes were grouped into three functional categories: adaptive/remodeling, inflammatory/immune-adhesive, and stress/damage response. Both angiogenic states engaged all three categories, but with different relative contributions. The 25 dynes/cm^2^ at 100 kPa state showed stronger expression for inflammatory and stress-associated genes, including cytokine- and immune-related genes (*e.g.*, *SPN*, *LCK*, *IL11*, *CSF3*) together with stress and damage response genes (*e.g.*, *NGFR*, *SAT1*, *SLC3A2*, *SQSTM1*, *BAK1*). In contrast, the 40 dynes/cm² at 10 kPa state showed relatively higher expression of adaptive and remodeling-associated genes, including *KLF2*, *ANGPT1*, *ITGB4*, *ADORA2A*, and *PRKAA2*, alongside genes involved in metabolic adaptation and structural remodeling. Although inflammatory and stress-response genes were also expressed under this condition (*e.g.*, *IL6*, *CXCL1*, *VCAM1*, *IL1A*, *CCL2*; *SEPTIN4*, *CD82*, *ACSL5*), they contributed less predominantly to the overall transcriptional program.

Therefore, whereas the 25 dynes/cm² at 100 kPa combination favored inflammatory and stress-associated signaling, the 40 dynes/cm^2^ at 10 kPa environments preferentially engaged adaptive and remodeling-associated pathways. Together, these findings extend the IPA predictions on SFN shifting the SS threshold for angiogenic activation, by showing that angiogenic activated states can be achieved through different transcriptional routes, involving different pathway balances depending on the experienced SS-SFN combination.

### 3.5 Transcriptional signatures recapitulate the bell-shaped SFN dependence of endothelial migration

Together, the IPA and KEGG analyses identified endothelial activation states associated with vascular remodeling under specific SS-SFN combinations. Because endothelial migration is a central component of vascular remodeling, we next examined whether migration-associated transcriptomic predictions were consistent with experimentally observed migration behavior. We previously showed that, on glass, the migration speed of ECs under flow peaks at 1 dyne/cm^2^ SS.^26^ Unpublished historical data from our group further showed that, under 1 dyne/cm^2^ SS, migration rates across substrates spanning 0.5-35 kPa SFN follow a bell-shaped profile across SFN, peaking at 4.5 kPa (Figure 5A).

**Figure 5:**
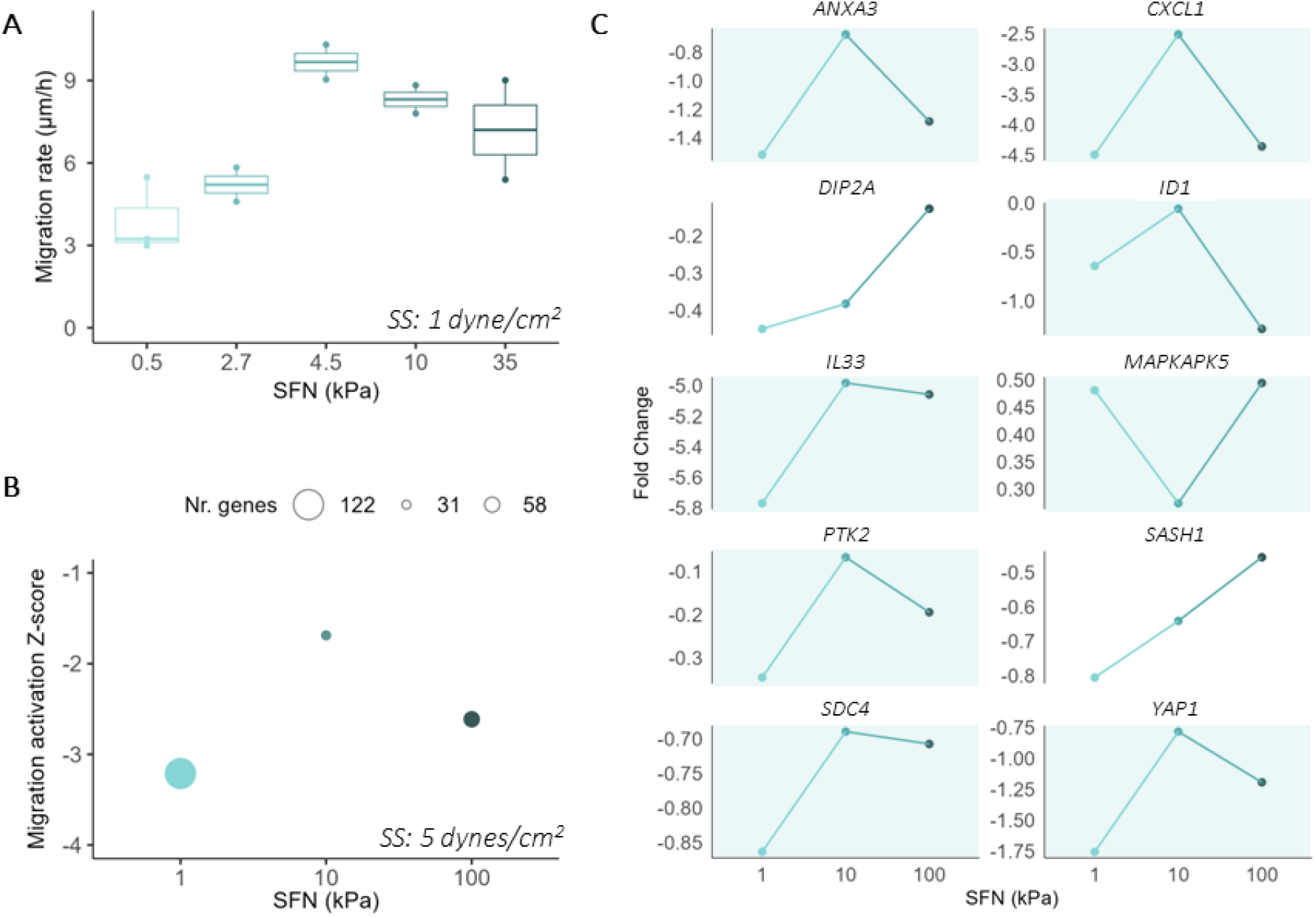
Correlation between transcriptomic predictions and *in vitro* data for EC migration under combined SS and SFN conditions. A, Migration rate of HUVECs cultured under 1 dyne/cm^2^ SS, and at 5 different levels of substrate SFN (0.5 – 35 kPa). B, IPA Z-scores for the migration of vascular ECs, based on gene fold changes at 5 dynes/cm^2^ SS vs. the reference condition (static SS at 100 kPa SFN) and at 3 levels of SFN (1, 10, 100 kPa). C, Gene expression fold changes for the genes in the INT list that were involved in EC migration. Turquoise backgrounds indicate genes with expression patterns matching *in vitro* EC migration rates and IPA predicted Z-scores for EC migration.

To determine whether our transcriptomic data reflected this behavior, we examined IPA activation Z-scores for endothelial migration at the lowest studied SS condition (5 dynes/cm^2^) across the three substrate SFNs (1, 10, 100 kPa). Relative to static, migration was predicted to be downregulated (negative Z-scores) at all three SFN levels, but the degree of inactivation followed the same bell-shaped dependence on SFN observed in the *in vitro* migration experiments, with the strongest predicted migration (or weakest inhibition) occurring at intermediate SFN (Figure 5B). The genes contributing to these predictions are listed in Table S10.

Among migration-associated genes identified by IPA under 5 dynes/cm^2^, ten were also present in the interaction gene set. Eight displayed expression patterns consistent with the SFN-dependent, bell-shaped migration profile observed experimentally (Figure 5C). Seven genes (*ANXA3*, *CXCL1*, *ID1*, *IL33*, *PTK2*, *SDC4* and *YAP1*) followed the same pattern direction, whereas *MAPKAPK5*, which has been reported to either promote or inhibit migration depending on cellular context,^36^ exhibited the inverted trend. Together, these findings indicate that migration-associated transcriptional signatures identified under low-SS reflect SFN-dependent endothelial migratory behavior observed *in vitro*.

### 3.6 YAP1 as a candidate integrator of shear stress and substrate stiffness cues

YAP1, together with its cofactor TAZ (*WWTR1*), is a central regulator of endothelial mechanotransduction and angiogenesis.^37^ Its activity is classically controlled by the Hippo signaling pathway.^38^ When Hippo signaling is inactive, YAP1/TAZ translocate to the nucleus, associate with TEAD transcription factors, and promote the expression of genes involved in proliferation and migration.^37,39^ Several observations from our transcriptomic analyses suggested that YAP1 may contribute to the integration of SS and substrate SFN. Both *YAP1* and its transcriptional partner *TEAD1* were identified among the 603 interaction-associated genes. IPA predicted *YAP1* as a regulator of proliferation, migration, and angiogenesis (Figure S5), and several established YAP1 target genes identified from ChIP-X Enrichment Analysis (ChEA) datasets,^40^ (including *FOXF1*, *PEAK3*, *PTX3*, *SEMA6A*, *SYNGR2*, or *TRAFD1*) were also present within the interaction gene set (Table S11). Together, these observations suggested that YAP1 activity may be influenced by the combined effects of SS and substrate SFN.

To investigate this possibility, we quantified active YAP1 nuclear localization across all SS-SFN conditions as a proxy for YAP1 transcriptional activity (Figure 6A-C). YAP1 nuclear localization (nuclear relative to whole-cell fluorescence intensities [N/C]) was markedly reduced on the softest substrate (1 kPa) compared with the stiffer substrates (mean N/C ratio 0.69-1.14 on 1 kPa vs. 1.57-2.27 10 and 100 kPa; *p* ≤ 3.7 x 10^-10^), and exhibited a significant SSxSFN interaction across conditions (two-way ANOVA, *p* = 2.34 x 10^-6^). In contrast, YAP1 remained predominantly nuclear across all SS conditions on the stiffer substrates, indicating reduced sensitivity to changes in SS.

**Figure 6:**
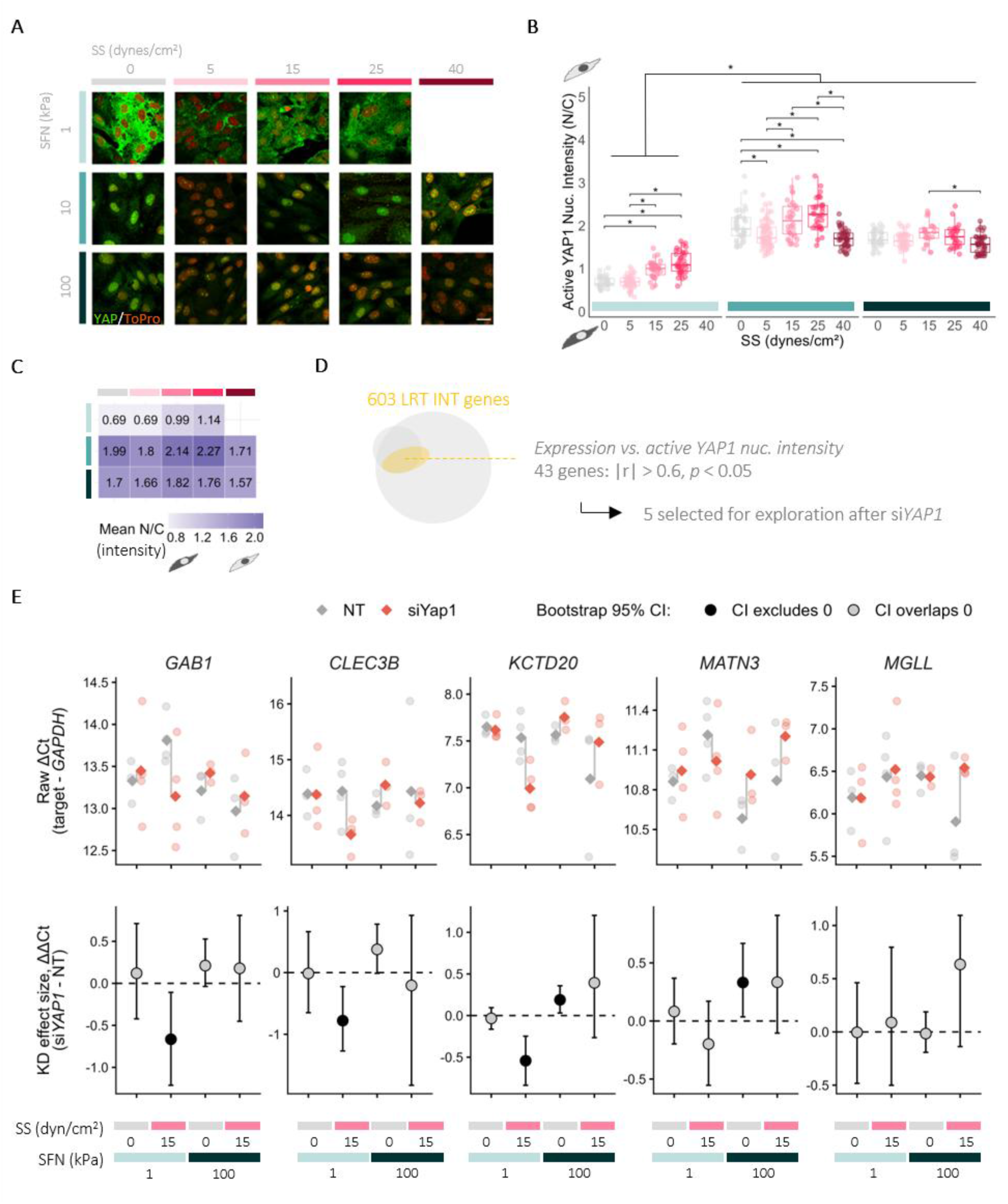
Context-dependent YAP1 activity across combined shear stress (SS) and substrate stiffness (SFN) cues. A, Representative immunofluorescence images of active YAP1 (green) and nuclei (TO-PRO, red). Scale bar = 20 µm. B, Relative active YAP1 nuclear localization (nuclear/whole cell intensities; N/C). Boxplots display the median, IQRs, and 1.5 x IQR whiskers (n ≥ 23 cells per condition). Asterisks indicate significant pairwise differences (two-way ANOVA with Tukey’s HSD, *p*-adj < 0.03). C, Heatmap of mean YAP1 nuclear localization across SS-SFN conditions. D, Correlation of YAP1 nuclear localization with the 603 interaction-associated genes identified by LRT (Supplementary Data 5). Five candidate genes were selected for exploratory analysis. E, Exploratory qPCR assessment following *YAP1* knockdown across four representative mechanical conditions. Upper panels show raw ΔCt values for individual biological replicates (n = 3-4; Supplementary Data 5), with mean values indicated by diamonds and gray connectors linking NT and si*YAP1* group means. Lower panels show the estimated knockdown effect size (ΔΔCt = si*YAP1* − NT) with 95% bootstrap confidence intervals (Supplementary Data 5).

To explore whether YAP1 was associated with interaction-responsive transcription, we correlated YAP1 nuclear localization with the expression profiles of the 603 interaction-associated genes across conditions. Forty-three genes significantly covaried with YAP1 nuclear localization (|r| > 0.6, Pearson *p* < 0.05), including 12 positive and 31 negative correlations (Figure 6D and Supplementary Data 5).

As an exploratory assessment of the contribution of YAP1 to interaction-responsive transcription, ECs were transfected with *YAP1* siRNA (80.4 ± 2.72% knockdown) and exposed to four representative SS-SFN conditions (1 or 100 kPa SFN under static or 15 dynes/cm^2^ SS). Expression of five candidate genes identified from the correlation analysis was then quantified as proof of principle (Figure 6E and Supplementary Data 5). The transcriptional effect of *YAP1* depletion was estimated as ΔΔCt together with its 95% confidence interval (CI); intervals excluding zero indicate a statistically supported effect under the corresponding mechanical condition. Most gene-condition combinations showed CIs overlapping zero. Nevertheless, four of the five candidate genes (*GAB1*, *CLEC3B*, *KCTD20*, and *MATN3*) exhibited at least one mechanical condition with a CI excluding zero, whereas *MGLL* showed only a comparatively large estimated effect despite its CI overlapping zero. Notably, the responsive mechanical condition differed among genes, and both the magnitude and direction of the transcriptional response varied across mechanical contexts, indicating that the consequences of *YAP1* depletion are mechanically context dependent.

Together with the interaction-dependent changes in YAP1 nuclear localization and its association with interaction-responsive gene expression, these proof-of-principle experiments support a role for YAP1 in integrating SS and substrate SFN in ECs.

## 4. Discussion

Although SS and SFN are two mechanical cues that coexist in blood vessels,^21,22^ their combined effects on endothelial transcription and behavior remain largely unexplored. Using a factorial RNA-Seq design across 14 mechanical conditions, coupled with a likelihood ratio-based analysis,^25^ we quantified the individual contributions of SS, SFN, and their interaction to endothelial gene expression. This approach identified 603 genes whose expression depended on the interaction between SS and SFN. By sampling multiple SS and SFN levels, our design revealed non-linear expression patterns and allowed us to distinguish shifts in the SS threshold for endothelial transcriptional activation from simple differences in response magnitude. Such behavior cannot be resolved reliably using conventional binary high-versus-low designs, which may overlook both threshold shifts and non-monotonic responses occurring at intermediate SS levels.

Consistent with their primary physiological role in sensing and responding to hemodynamic forces, ECs exhibited substantially greater transcriptional sensitivity to SS than to substrate SFN. This broader sensitivity to SS may reflect the continuous fluctuations in flow associated with cardiac pulsatility, exercise, vascular branching, and pathological disturbances, whereas substantial changes in substrate SFN generally develop over longer timescales during aging, fibrosis, or tissue remodeling. Nevertheless, SFN was biologically active and modified the expression of more than 12% of SS-responsive genes through additive and interaction-dependent effects. These findings support a hierarchical model of endothelial mechanosensing in which SS defines the dominant transcriptional landscape, while substrate SFN modulates how ECs interpret flow. This behavior is consistent with previous work showing that subendothelial SFN substantially alters endothelial mechanics and traction force generation while exerting only modest effects on the transcriptome.^41^ One possible explanation is that SS and SFN converge, at least in part, on shared force-transmission machinery influencing traction forces through apical cytoskeletal tension and basal adhesion mechanics, respectively. Both inputs ultimately modulate cytoskeletal tension and traction generation, which can propagate matrix deformations over distances exceeding 30 µm,^42^ depending on substrate SFN and viscosity –a length scale relevant in perivascular fibrosis and ECM remodeling.

Endothelial responses across increasing SS levels were frequently non-linear. Although biphasic responses to SS have been reported for individual genes and proteins (e.g., *MCP-1* expression and Smad2/3 activation),^43,44^ previous studies have largely examined discrete flow regimes^45^ or temporal dynamics,^46^ leaving transcriptome-wide responses across SS magnitude insufficiently characterized. Importantly, non-linear responses provide the basis for thresholded behavior, allowing ECs to discriminate different flow regimes and selectively engage distinct transcriptional programs rather than responding proportionally to increasing force. Accordingly, low SS is generally associated with pro-inflammatory and atheroprone phenotypes, whereas intermediate SS promotes endothelial quiescence and vascular homeostasis,^47^ while higher SS levels engage adaptive stress-responsive processes. The highest SS examined here (40 dynes/cm^2^) falls within the upper physiological range reported for some arterial and valvular environments, and although it likely exceeds that experienced by HUVECs, is well below the extreme pathological SS (>100 dynes/cm^2^) observed in highly stenosed arteries.^48^ Collectively, these observations suggest that the non-linear transcriptional patterns provide the mechanistic basis for endothelial flow set points, allowing transitions between transcriptional programs as mechanical environments change.

A central finding of this study was that substrate SFN shifted the SS threshold at which angiogenesis- and remodeling-associated transcriptional programs become activated. Both IPA-and KEGG-based analyses supported activation at 25 dynes/cm^2^ on 100 kPa substrates and at 40 dynes/cm^2^ on 10 kPa substrates, suggesting that this apparent set point is not fixed but depends on the mechanical properties of the substrate, with increased SFN generally shifting toward lower SS levels. This threshold behavior also helped explain the organization of the global transcriptomic landscape. ECs exposed to low SS (5 dynes/cm^2^) formed a distinct transcriptional group (cluster 1), consistent with the differentiated phenotype associated with low-flow environments. In contrast, higher SS conditions (15, 25, and 40 dynes/cm^2^) occupied a broader and more heterogeneous UMAP region that could not be resolved by SS or SFN magnitude alone (clusters 2 and 3). Within this space, angiogenesis-associated activation emerged as a key organizing feature, segregating samples into angiogenesis-inactive (cluster 2) and angiogenesis-active (cluster 3) states. Thus, the modulation of the SS threshold for angiogenic activation by substrate SFN provides a unifying explanation for the transcriptional heterogeneity observed under elevated SS.

Mechanical context influenced not only when angiogenesis-associated programs became activated, but also the composition of the resulting transcriptional profile. The conditions of 25 dynes/cm^2^ on 100 kPa and 40 dynes/cm^2^ on 10 kPa both shared core features of endothelial activation –including NF-κB signaling, which has a key role in initial inflammatory activation.

However, the former showed a stronger relative contribution from inflammatory, immune- adhesive, and stress-related programs, whereas the latter displayed a greater contribution from adaptive and remodeling-associated processes. These results suggest that different mechanical routes converge on broadly related endothelial activation programs through context-dependent reweighting of mostly shared transcriptional modules. Accordingly, while SFN shifts the SS set point that determines when inflammation and further remodeling begin, the specific SS-SFN combination (or mechanical route) that activated it influences which components of the remodeling response dominate. Further studies, however, should determine the biological impact that this mechanically driven divergence between activated endothelial states has on EC behavior.

The non-monotonic responses observed across the mechanical landscape may be interpreted within frameworks of force transmission such as the integrin-based molecular clutch model.^49,50^ Efficient mechanotransduction depends on the dynamic coupling of integrins, focal adhesions, and the actomyosin cytoskeleton, and this coupling varies with substrate mechanics.^50^ Changes in SFN may therefore alter the transmission of flow-induced cytoskeletal forces and shift the SS required to engage downstream signaling. This would help explain why angiogenesis-associated activation emerged at 25 dynes/cm^2^ on 100 kPa substrates but was attenuated at 40 dynes/cm^2^, as well as why migration exhibited a biphasic dependence on SFN. However, because adhesion dynamics and traction forces were not measured directly, this interpretation remains a hypothesis for future investigation.

YAP1 provided an additional mechanistic perspective on SS-SFN integration. *YAP1* and *TEAD1* belonged to the interaction-responsive gene set, multiple established YAP1 targets displayed interaction-dependent expression, and YAP1 nuclear localization showed a significant SS-SFN interaction. On soft substrates, YAP1 was relatively excluded from the nucleus but became more nuclear with increasing SS, whereas on stiffer substrates YAP1 remained predominantly nuclear and was less responsive to changes in SS. Previous studies have shown that YAP1 integrates both flow and substrate mechanics, with flow direction regulating YAP1 nuclear localization during cardiac valve development and pathological substrate stiffening attenuating flow-induced YAP1 nuclear exclusion.^51,52^ Our data extend these observations by showing that YAP1 responsiveness depends on the specific combination of SS and substrate SFN across a physiologically relevant mechanical landscape.

Exploratory *YAP1* knockdown experiments further indicated that the transcriptional consequences of YAP1 perturbation varied by gene and mechanical condition. The direction and magnitude of the observed effects were not consistent across the SS-SFN landscape, suggesting that YAP1 is unlikely to control interaction-responsive transcription alone. Rather, YAP1 may function as one component of a broader network emerging from the coordinated activity of multiple mechanosensitive pathways. The present data therefore support a context-dependent association between YAP1 and SS-SFN integration but do not establish YAP1 as the causal mediator of the threshold shift. More comprehensive perturbation studies will be needed to define its contribution and interaction with other mechanosensitive regulators.

These findings have important implications for experimental vascular models. Most conventional flow systems expose ECs cultured on glass or plastic to a selected SS regime, thereby combining flow with a substrate many orders of magnitude stiffer than most native tissues. Our results indicate that the transcriptional state elicited by a particular SS cannot be extrapolated across substrates of different SFNs. Models that reproduce flow while neglecting physiologically relevant matrix mechanics may therefore misrepresent endothelial sensitivity and remodeling behavior.

Incorporating both cues should improve the interpretation of vascular disease models, organ-on-chip platforms, and tissue-engineered constructs, particularly in settings involving fibrosis, aging, tumor-associated stiffening, or biomaterial implantation.

Several limitations should be acknowledged. HUVECs were used as a reproducible endothelial model, but ECs exhibit pronounced heterogeneity across vascular beds, organs, and arterial-venous hierarchies,^53^ including tissue-specific mechanobiological responses,^54^ and the transcriptional responses observed here may not fully capture the behavior of specialized subtypes. In addition, ECs mechanotransduce differently on collagen than on fibronectin^55^ and therefore different sets of genes may arise using fibronectin-functionalized hydrogels. Moreover, we used purely elastic substrates, whereas native tissues are viscoelastic and dissipate forces over time. Finally, we applied steady laminar SS without physiological pulsatility, multidirectional flow, and circumferential strain. These additional mechanical cues may further modulate the endothelial integration of SS and SFN.

## Conclusions

To our knowledge, this study provides the first systematic analysis of how SS and substrate SFN jointly shape endothelial transcriptomic responses across a graded mechanical landscape. Using a factorial transcriptomic approach, we distinguished genes modulated by SS or SFN alone from those co-modulated by their interaction, revealing that substrate SFN shifts the shear stress threshold at which angiogenesis-associated transcriptional programs become activated.

Moreover, the qualitative balance of the resulting transcriptionally activated response depended on the combined mechanical context, reflecting different balances of adaptive, inflammatory, and stress-responsive programs. Together, these findings support a hierarchical model of endothelial mechanosensing in which SS defines the dominant transcriptional landscape, while substrate SFN tunes the endothelial sensitivity to flow and the nature of the resulting remodeling profile.

Incorporating both mechanical cues into experimental vascular models will be important for accurately reproducing endothelial physiology and interpreting vascular mechanobiology.

## Funding

This project was supported by funding from

European Union’s Horizon 2020 research and innovation programme under the Marie

Skłodowska-Curie grant agreement No 101025264, H2020-MSCA-IF-2020, ECSiTe (LGR)

KU Leuven internal funding IDN 19/031 (EAVJ, AL, HVO)

Fonds Wetenschappelijk Onderzoek, FWO funding, G0GE323N (EAVJ)

## Acknowledgments

We thank Dr. R. Sinha and Dr. S. Simmonds for their support in the design of the setup to prepare hydrogels at the different stiffnesses; M. Brusselmans for his biostatistics support on the design of the factorial RNA-Seq analysis; Dr. A. Cortés-Calabuig, Dr. F. Ver Donck, Dr. S. Ekhteraeitousi and Dr. J. Mathys for their discussions on the RNA-Seq analysis; the KU Leuven Genomics Core, and Dr. S. Plaisance for their support with the IPA software; the members from the CMVB groups at KU Leuven for their feedback on the analysis and results.

## Author contributions

Study conception and experimental design: LGR, EAVJ

Feedback on experimental design: AL, HVO.

Methodology: LGR, HVO, EAVJ developed the flow setup to combine shear and stiffness, AT

developed the migration flow setup with the cell tracking system.

Investigation: LGR performed all experiments except for *in vitro* EC migration, by AT. FB prepared the hydrogels for the knock down experiment.

Data curation: LGR produced and maintained all metadata regarding samples and wrote the R codes.

Analysis – RNA-Seq: LGR, EAVJ.

Analysis – phenotypical changes: WG, LGR

Results interpretation: LGR, AL, HVO, EAVJ

Visualization: LGR, EAVJ.

Supervision: EAVJ.

Writing – original draft: LGR, EAVJ.

Writing – review & editing: LGR, AT, WG, FB, AL, HVO, EAVJ.

## Supplementary Materials

Supplementary Figures S1 – S7

Supplementary Tables S1 – S11

Supplementary Movie 1

Supplementary Data 1 – 5 (available upon publication)

Supplementary Methods

References in Supplementary Materials

