## Supplementary Materials for "Matrix stiffness shifts the endothelial shear stress set point for angiogenic activation"

**Short title:** Integration of shear stress and stiffness in remodeling

*Laia Gifre-Renom, Ashkan Tabibian, Wolfgang Giese, Femke Bellen, Aernout Luttun, Hans Van Oosterwyck, Elizabeth A.V. Jones**

**This PDF file includes:**

Supplementary Figures S1 – S7

Supplementary Tables S1 – S11

Supplementary Movie 1

Supplementary Data 1 – 5 (available after publication in journal)

Supplementary Methods

References in Supplementary Materials

### Supplementary Figures

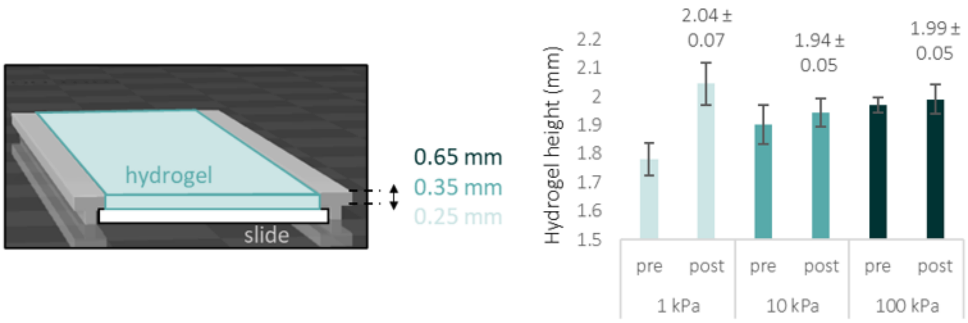


**Figure S1**. T-shaped spacers were designed (illustrated in gray) to ensure that the different polyacrylamide swelling in 1, 10 and 100 kPa hydrogels would result in a 2 mm total height (including 1 mm slide) after overnight incubations in DPBS. Since the different stiffness (SFN) hydrogels swelled differently, this was compensated by designing the top part of the spacers (horizontal part in the T shape) to measure exactly 0.65, 0.35, and 0.25 mm for 100, 10, and 1 kPa hydrogels, respectively. Three measurements (top, mid, bottom part) across three slides (containing the corresponding hydrogel) were performed per each SFN at each timepoint. Mean and standard errors for the measurements are plotted per each SFN. The flow chambers for 1 kPa hydrogels were further adapted deepening them by 0.1 mm, since spacers thinner than 1.25 mm (0.25 for the hydrogel part) were not printable.

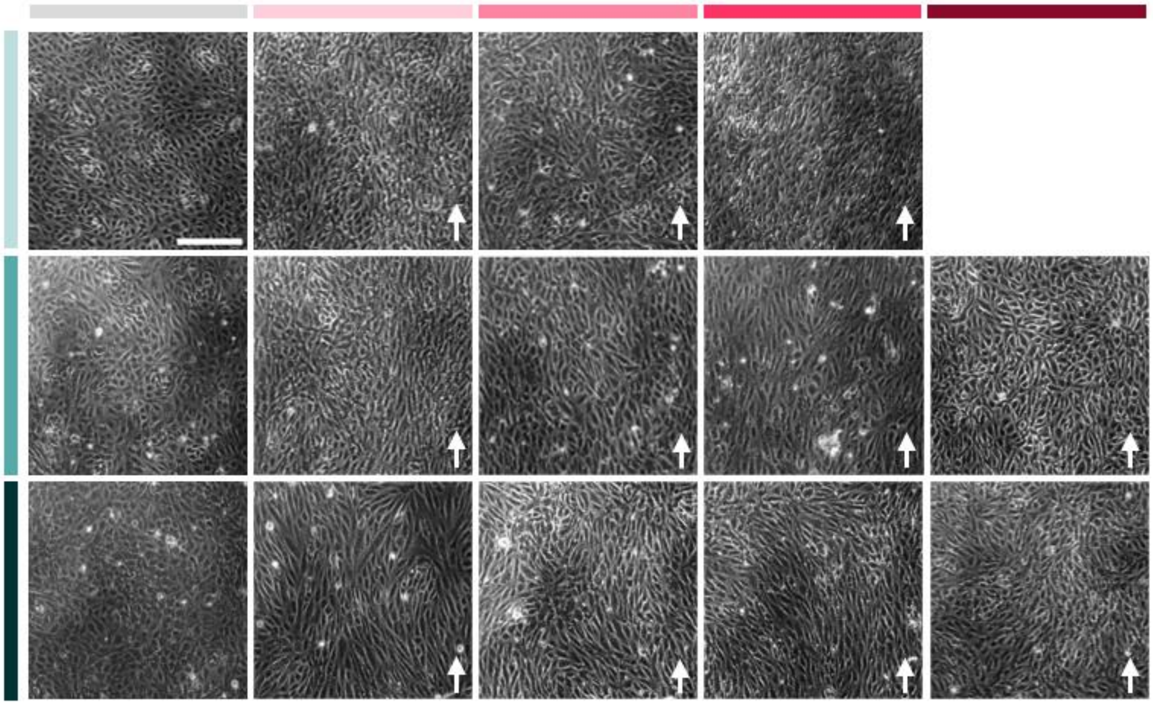

Figure S2. Alignment of HUVECs under shear stress (SS; 0, 5, 15, 25, 40 dynes/cm$^2$; gray for static, darkening pink palette for SS levels) at different stiffnesses (1, 10, 100 kPa; darkening turquoise palette). Scale bar indicates 200 µm and applies for all images. Arrows indicate the direction of flow.

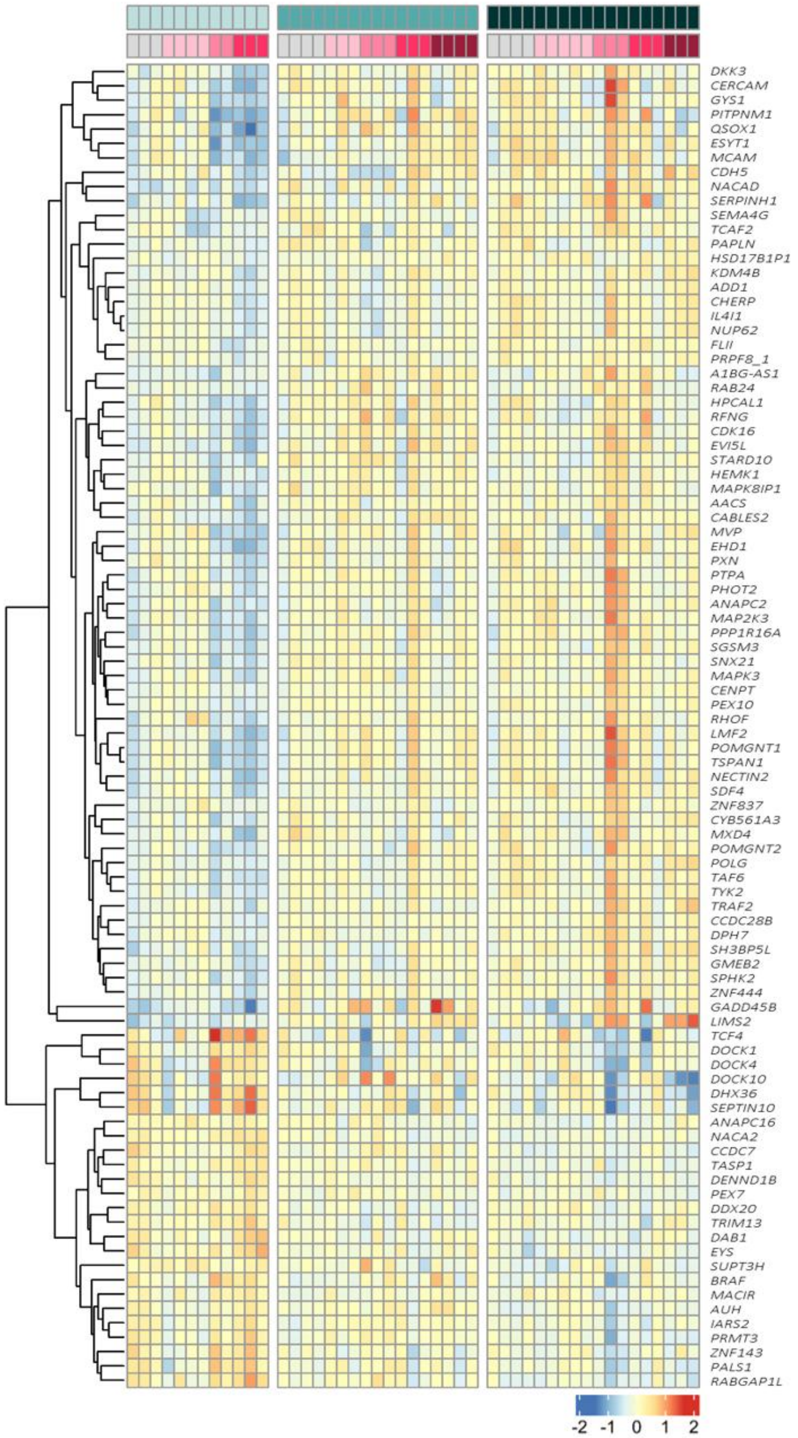


**Figure S3.** Heatmap showing the variation across conditions for genes related to stiffness (SFN). The selected genes are a combination between literature-based genes (*PXN, RHOF, RHOT2, CDH5, LIMS2, DOCK1, DOCK4, DOCK10*) and the 96 unique-SFN genes obtained by Likelihood ratio test (LRT) comparing the differential expression from a full model (containing shear stress or SS, SFN and interaction or INT terms) and a reduced model (where SFN terms were excluded; n = 3 to 5 samples per condition; n = 2 for 1 kPa at 15 dynes/cm$^2$; $p$-adj < 0.05).

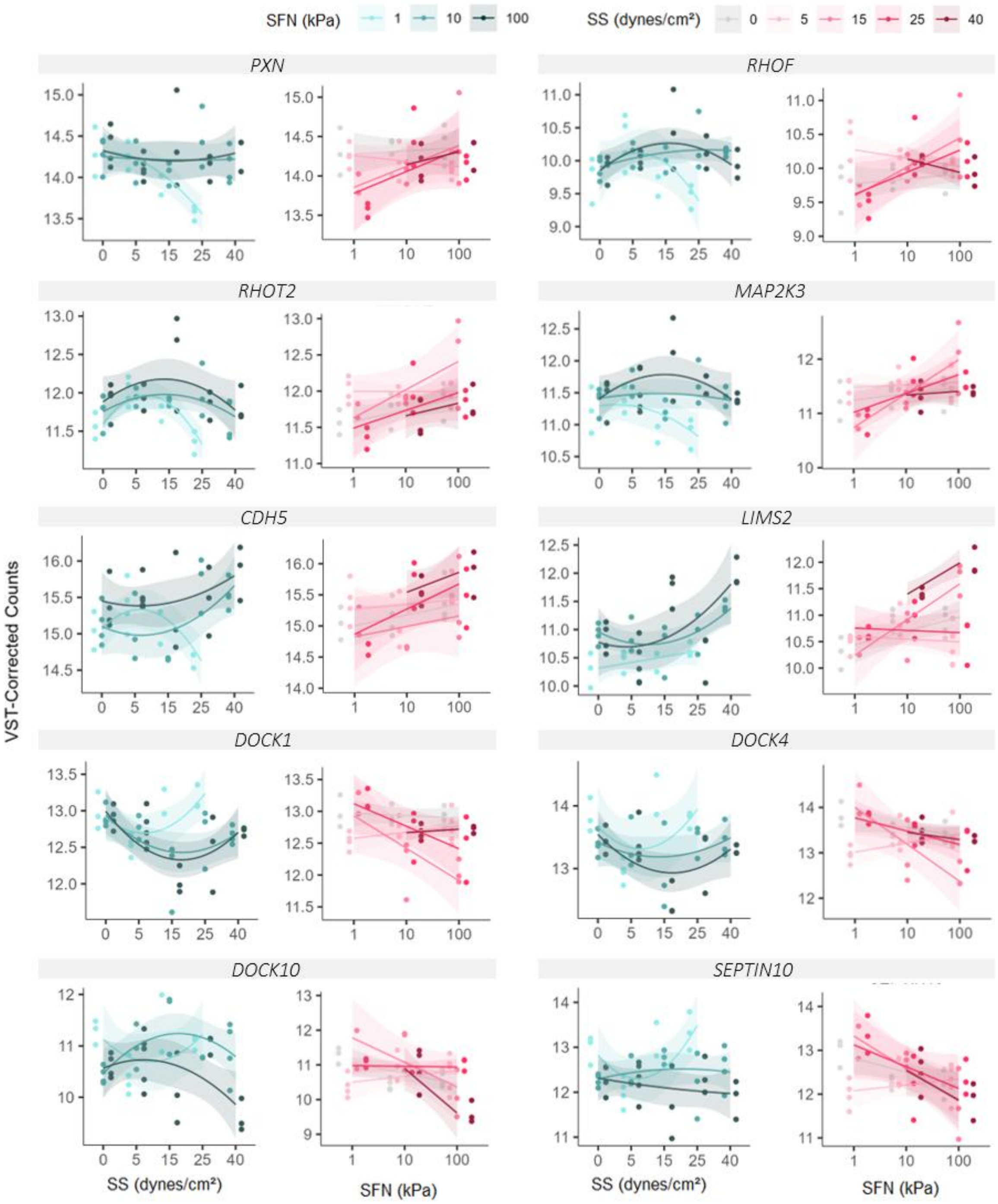


Figure S4. Variance Stabilizing Transformation (VST)-corrected gene count dynamics across conditions for genes with changes across substrate stiffness (SFN) (n = 3 to 5 samples per condition; n = 2 for 1 kPa at 15 dynes/cm$^2$). Left-handed plots, counts across the 5 levels of shear stress (SS), colored by SFN. Expression dynamics represented by quadratic regression curves ("lm" smoothing method with formula y ~ poly(x, 2)) for SFN across SS with standard error (SE) shades. Right-handed plots, counts across the 3 SFN levels, colored by SS. Expression dynamics represented by regression lines ("lm" smoothing method with formula y ~ x)) for SS across SFN with SE shades.

A

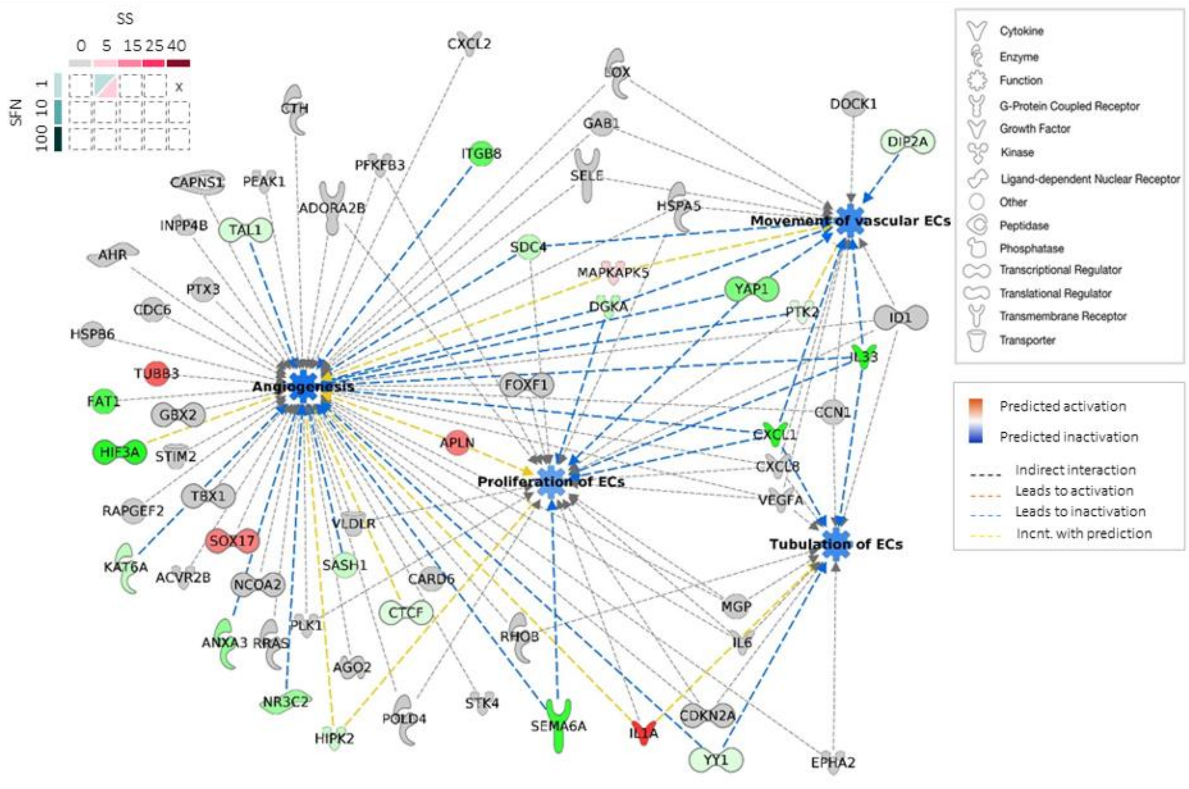


© 2000-2023 QIAGEN. All rights reserved.

B

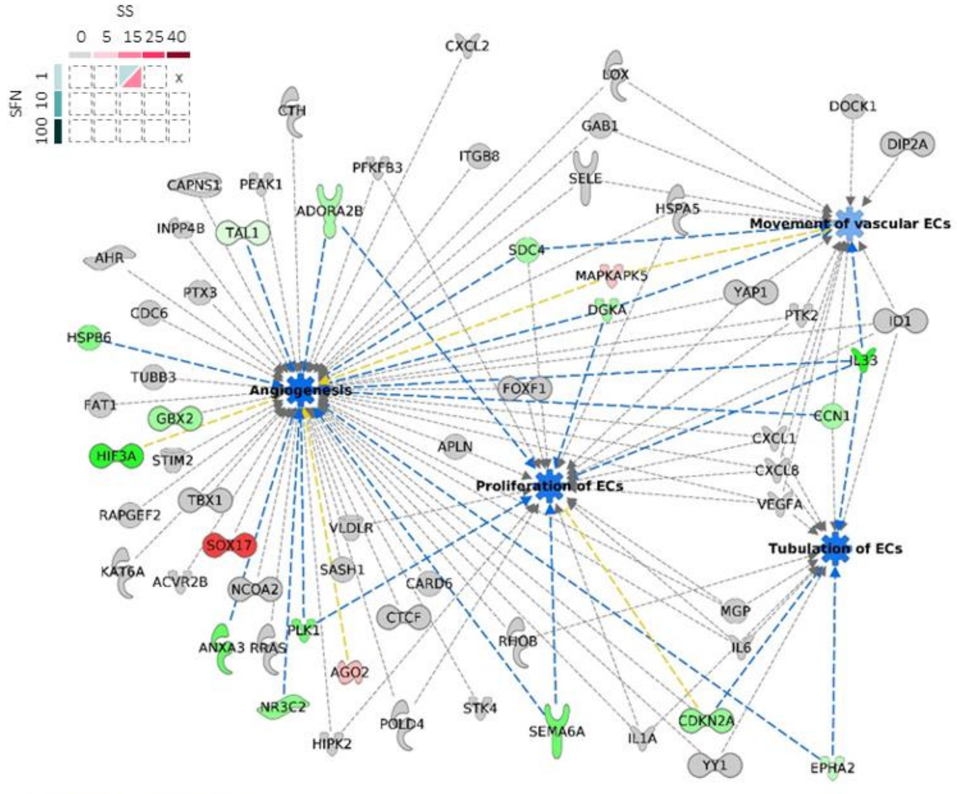


© 2000-2023 QIAGEN. All rights reserved.

C

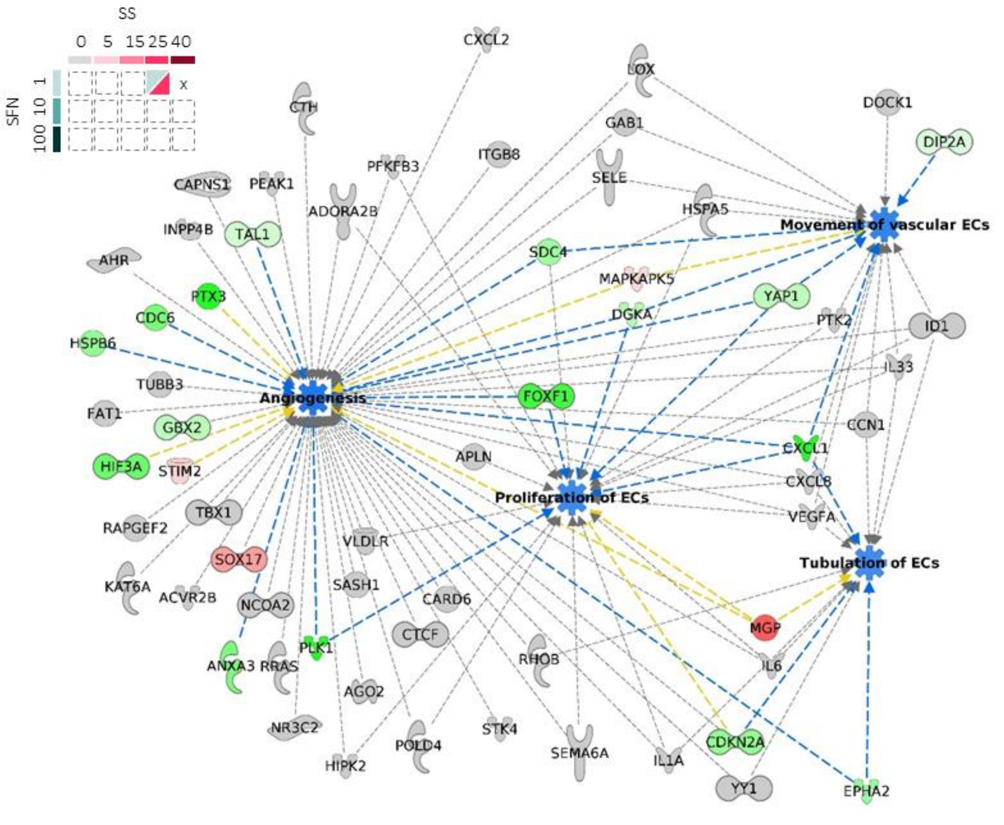


© 2000-2023 QIAGEN. All rights reserved.

D

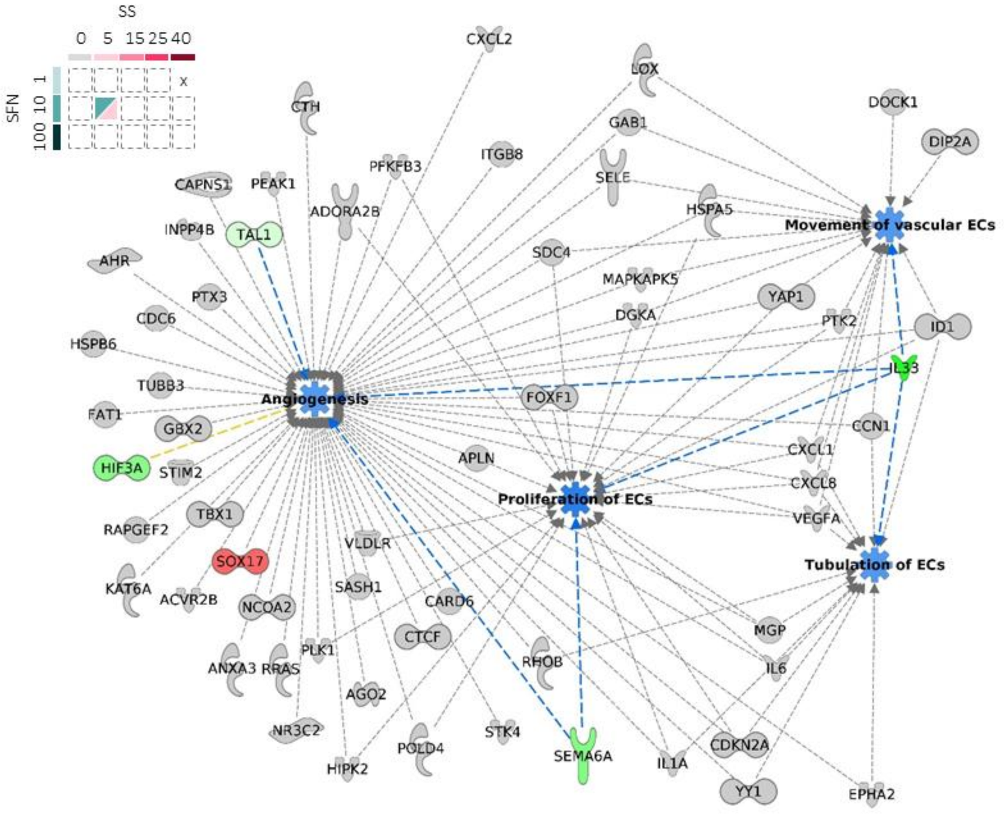


© 2000-2023 QIAGEN. All rights reserved.

E

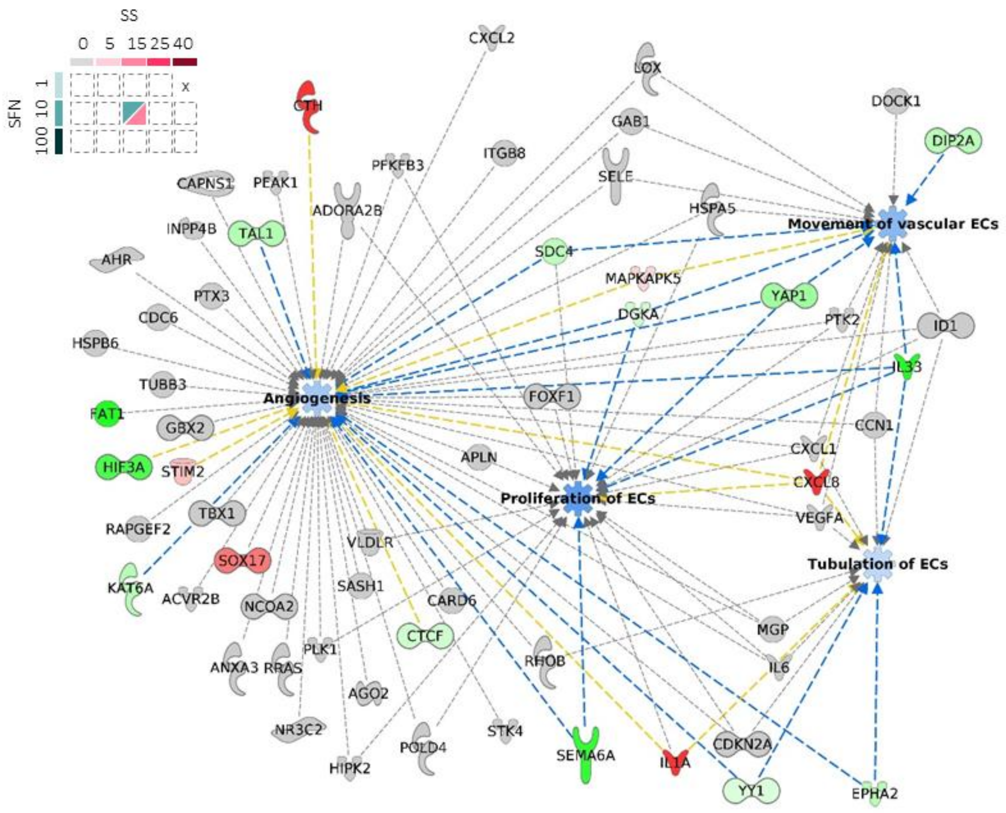


© 2000-2023 QIAGEN. All rights reserved.

F

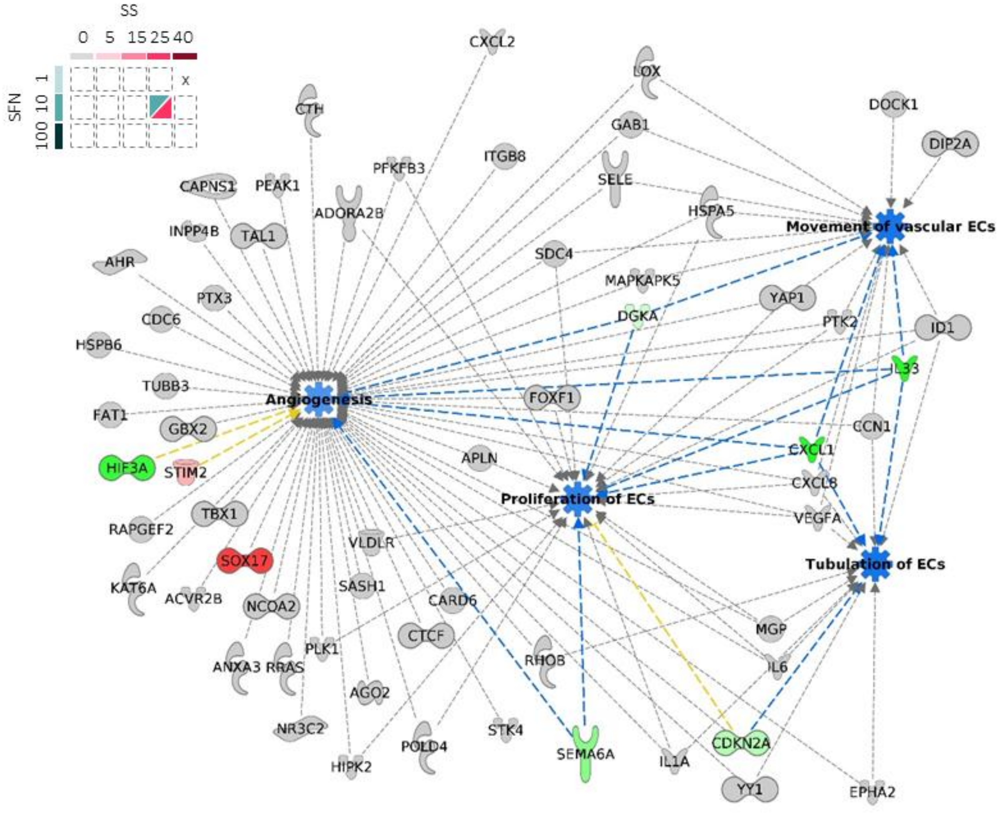


© 2000-2023 QIAGEN. All rights reserved.

G

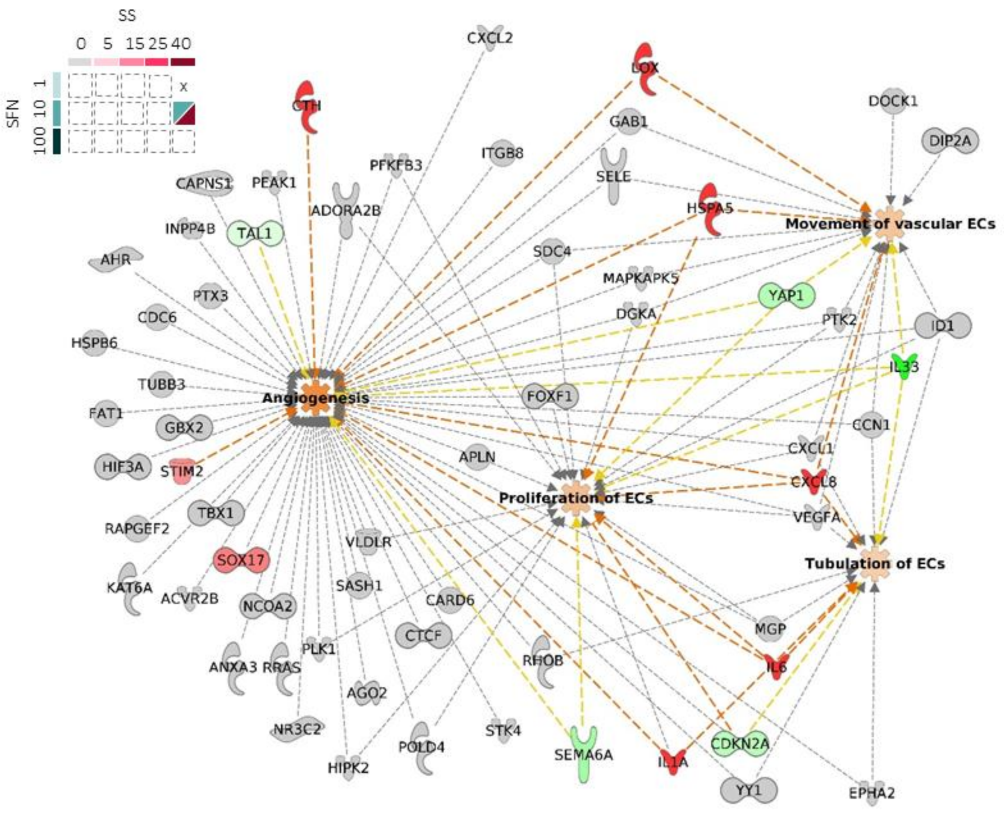


© 2000-2023 QIAGEN. All rights reserved.

H

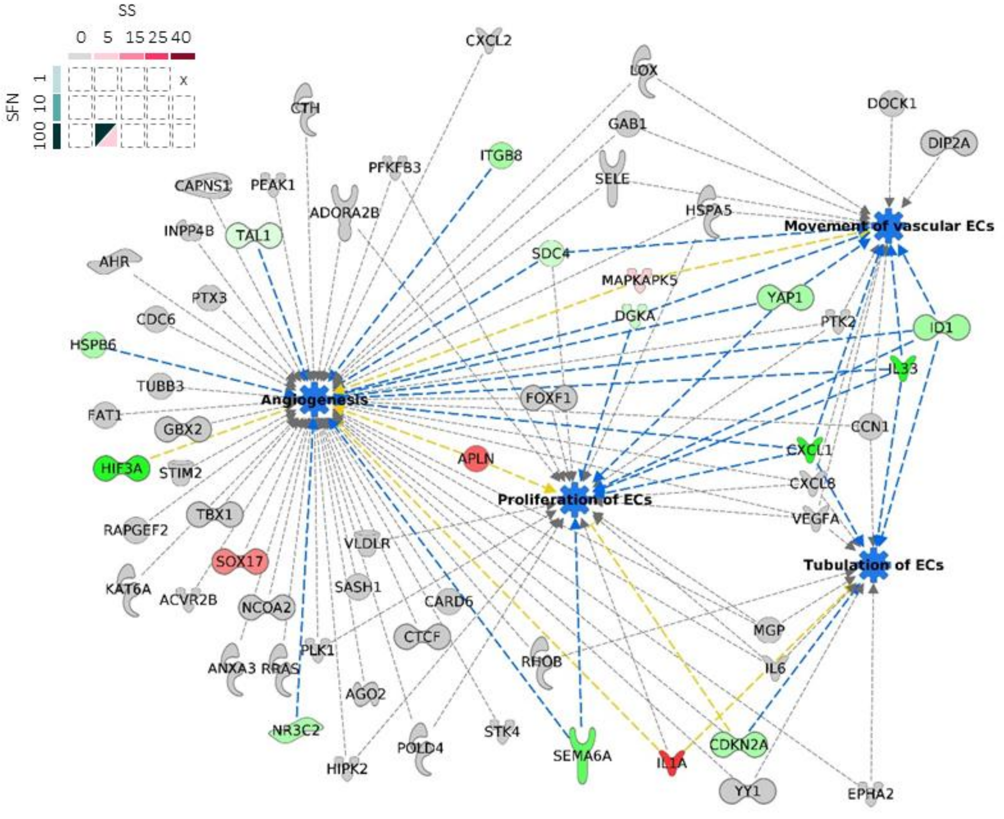


© 2000-2023 QIAGEN. All rights reserved.

I

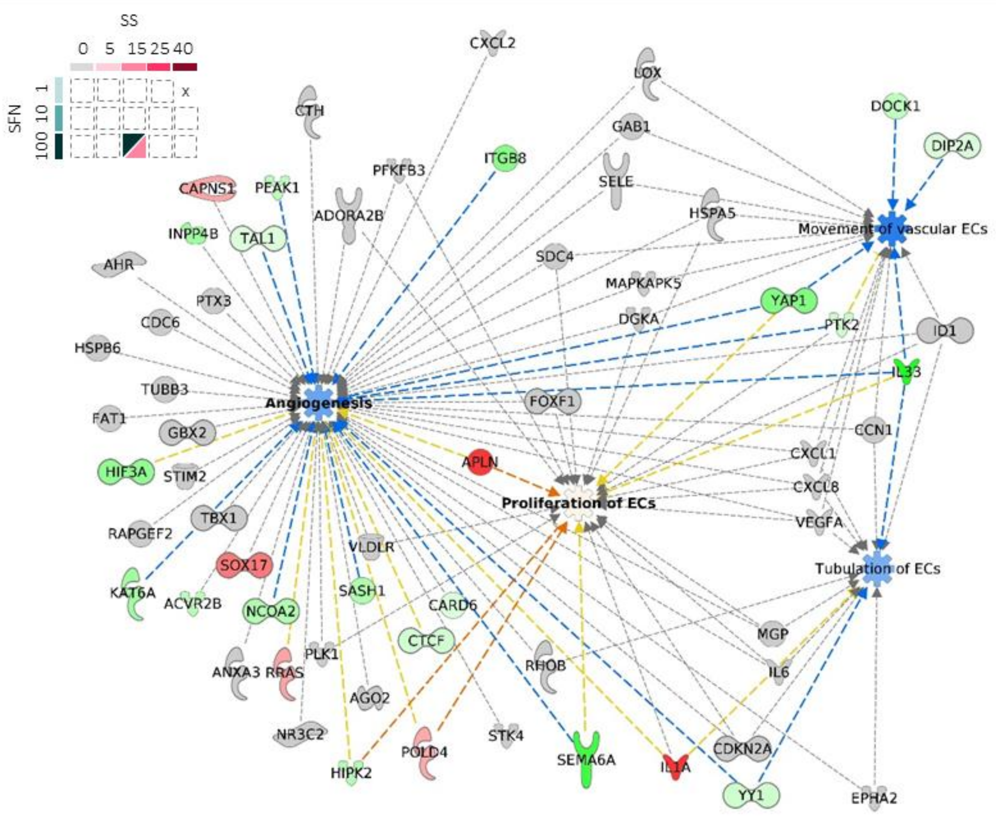


© 2000-2023 QIAGEN. All rights reserved.

J

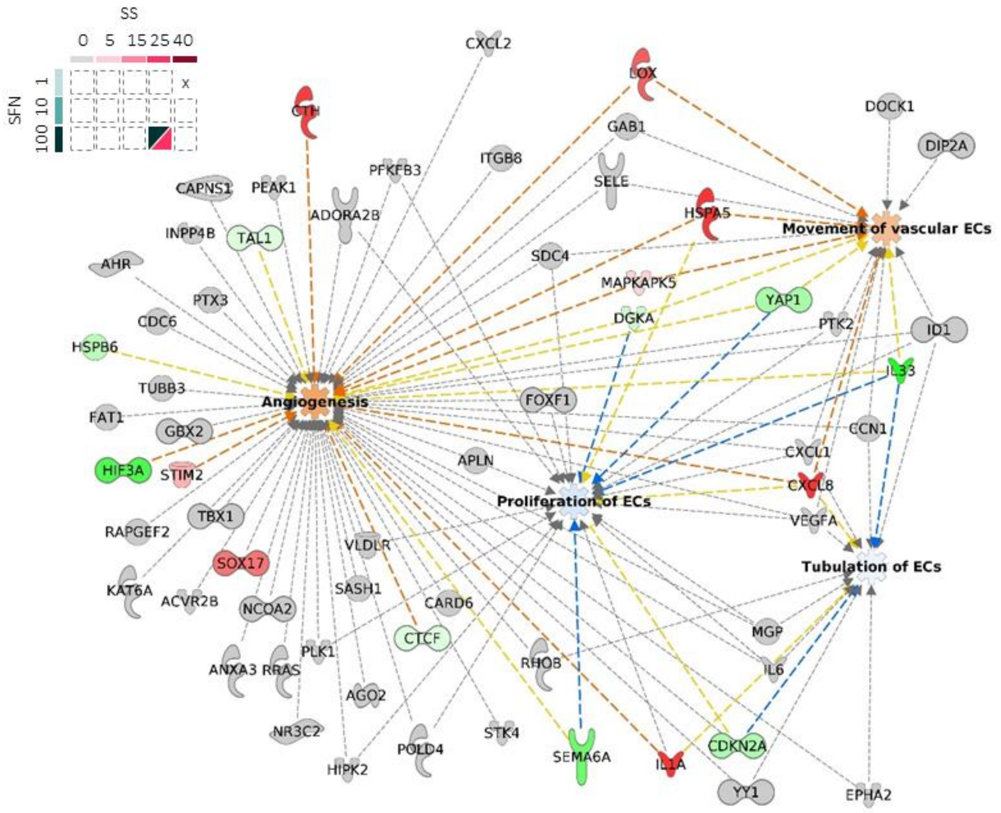


© 2000-2023 QIAGEN. All rights reserved.

K

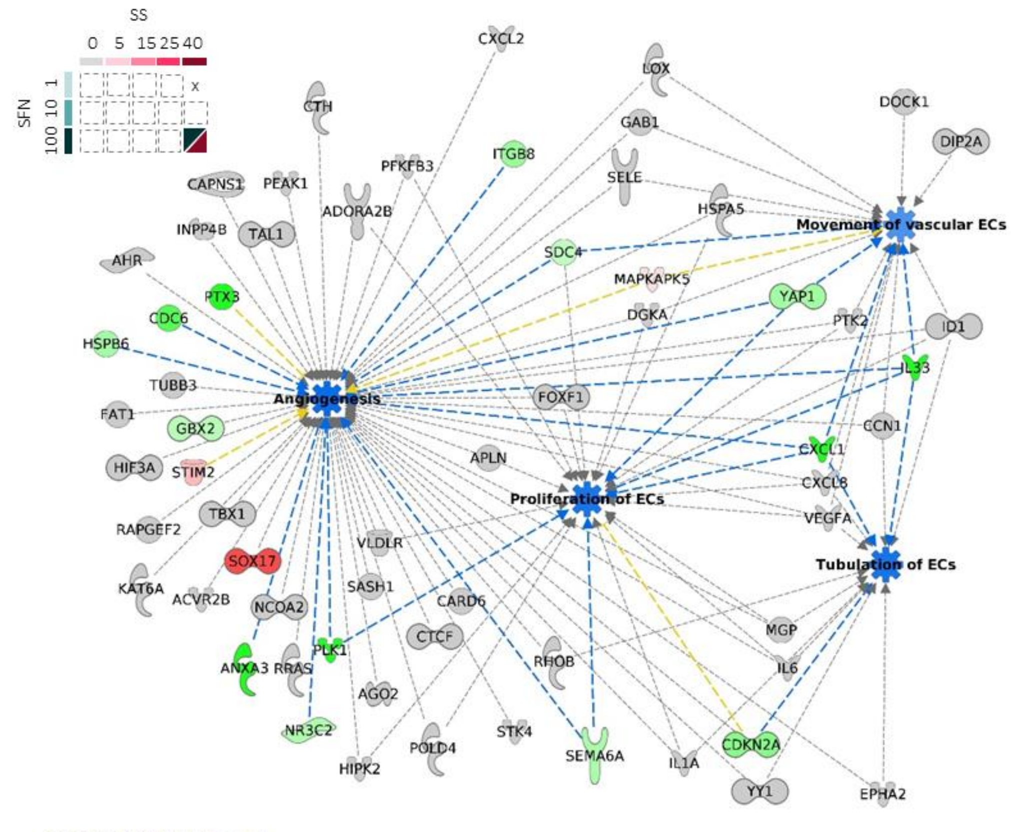


© 2000-2023 QIAGEN. All rights reserved.

**Figure S5.** Complete process networks across shear stress (SS) and stiffness (SFN) combined conditions, including the gene names and their fold change color code (green, up-regulated; red, down-regulated; adj-$p < 0.05$). Conditions are: 1 kPa at 5 dynes/cm$^2$ (**A**), 15 dynes/cm$^2$ (**B**), and 25 dynes/cm$^2$ (**C**); 10 kPa at 5 dynes/cm$^2$ (**D**), 15 dynes/cm$^2$ (**E**), and 25 dynes/cm$^2$ (**F**), 40 dynes/cm$^2$ (**G**); 100 kPa at 5 dynes/cm$^2$ (**H**), 15 dynes/cm$^2$ (**I**), and 25 dynes/cm$^2$ (**J**), 40 dynes/cm$^2$ (**K**). Positive Z-scores predict the activation of a process, given that they are obtained when enhancer genes are upregulated, or when inhibitor genes are downregulated, and *vice-versa* for negative Z-scores. A color-coded network was then obtained for each tested condition: genes were colored in red or green when these were observed up or downregulated (positive or negative fold changes, respectively; adj-$p < 0.05$); processes were colored orange or blue when these were predicted to be activated or inactivated (Z-scores > 2 or < -2, respectively; adj-$p < 0.05$). Indirect interactions between predicted processes and observed genes are represented as dashed lines. Yellow lines indicate inconsistent findings with the node predictions. An animated version of this network is available as *Supplementary Movie 1.*

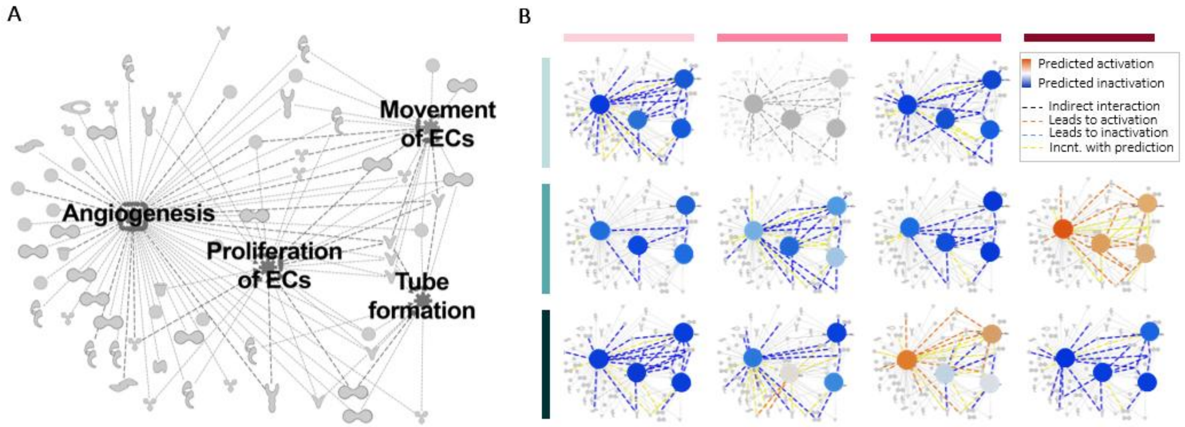


**Figure S6. A**, Summarized network for the 4 selected top "cardiovascular system development and disease" biological processes obtained by Ingenuity Pathway Analysis (IPA) from the 603 genes in the interaction LRT list; gene names and shape codes are provided in Supplementary Fig. S4. **B**, Full overview of IPA predicted in/activation state for cardiovascular processes across the 11 non-static conditions, calculated from the gene fold changes and their enhancer/inhibitor reported role in the process (Z-scores). Significant predictions are Z > 2 for activation (orange) and Z < -2 for inactivation (blue). Indirect interactions between predicted processes and observed genes are represented as dashed lines. Yellow lines indicate inconsistent findings with the node predictions. IPA predictions for condition 15 dynes/cm$^2$ SS at SFN 1 kPa were based on n = 2; depicted in gray.

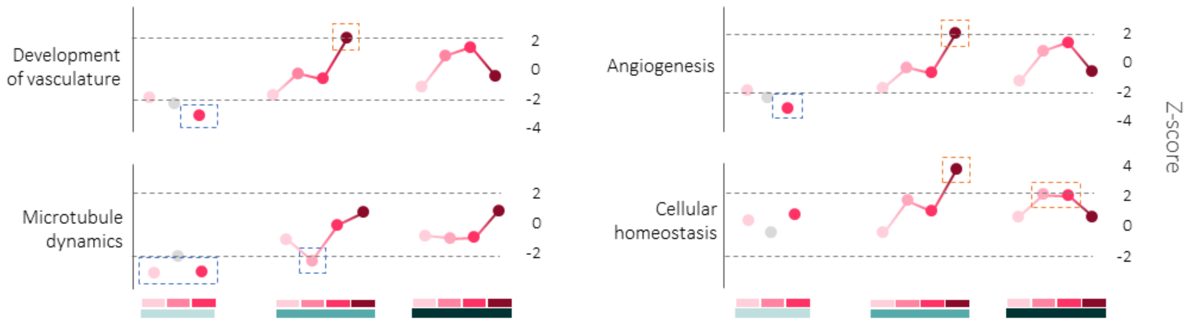


**Figure S7.** Representation of the IPA process activation Z-score dynamics across conditions for development of vasculature, angiogenesis, microtubule dynamics, and cellular homeostasis. Z-score points are colored by shear stress (SS) levels and grouped by stiffness (SFN) degree. Significant inactivation (blue) and activation predictions (orange) are depicted in dashed boxes ($p < 0.00002$). IPA predictions for condition 15 dynes/cm$^2$ SS at SFN 1 kPa were based on n = 2; depicted in gray.

### Supplementary Tables

**Table S1.** Poly-acrylamide proportions in hydrogels at different stiffnesses in RNA-Seq experiments.

| Stiffness | Acrylamide | Bis-Acrylamide | Reference |
|---|---|---|---|
| 1 kPa | 5 % | 0.03 % | 1 |
| 10 kPa | 10 % | 0.075 % | 2 |
| 100 kPa | 10 % | 0.4 % | 3 |

**Table S2.** Pump programs used to apply shear stress on HUVECs.

| Shear Stress | Ramp-up time | Initial flow rate | Final flow rate | Steady time |
|---|---|---|---|---|
| 5 dynes/cm$^2$ | - | 17.5 mL/min | 17.5 mL/min | 24 h |
| 15 dynes/cm$^2$ | 1 h | 35 mL/min | 52.85 mL/min | 23 h |
| 25 dynes/cm$^2$ | 3 h | 35 mL/min | 87.5 mL/min | 21 h |
| 40 dynes/cm$^2$ | 6 h | 35 mL/min | 140 mL/min | 18 h |

**Table S3.** Summary of the replicates for the 14 conditions.

| SFN | | | SS | | |
|---|---|---|---|---|---|
| | Static | 5 dynes/cm$^2$ | 15 dynes/cm$^2$ | 25 dynes/cm$^2$ | 40 dynes/cm$^2$ |
| Replicates in flow experiments | | | | | |
| 1 kPa | 3 | 4 | 3 | 4 | - |
| 10 kPa | 4 | 4 | 3 | 4 | 4 |
| 100 kPa | 5 | 5 | 3 | 4 | 3 |
| Replicates used for differential expression analysis | | | | | |
| 1 kPa | 3 | 4 | 2 | 3 | - |
| 10 kPa | 4 | 3 | 3 | 3 | 4 |
| 100 kPa | 4 | 5 | 3 | 3 | 3 |

**Table S4.** Quality control* on the RNA-Seq data and sample exclusion.

| | sample | total reads (Hisat2) | total assigned | unmap. | multi-mapping | no features | % assig. from tot. reads **(>70%)** | % unmap. | % multi-map. **(<2SD)** | % no feat. |
|---|---|---|---|---|---|---|---|---|---|---|
| 7 sample repeat | reS | 111255078 | 74607231 | 1347535 | 35095527 | 204785 | 67.1 | 1.2 | 31.5 | 0.2 |
| | reAT | 77566833 | 54752181 | 1027984 | 21565339 | 221329 | 70.6 | 1.3 | 27.8 | 0.3 |
| | reBW | 84994503 | 60681827 | 823969 | 23332663 | 156044 | 71.4 | 1.0 | 27.5 | 0.2 |
| | reAF | 79143326 | 58609498 | 743885 | 19584683 | 205260 | 74.1 | 0.9 | 24.7 | 0.3 |
| | reBX | 70385776 | 52863116 | 580743 | 16745451 | 196466 | 75.1 | 0.8 | 23.8 | 0.3 |
| | reBK | 77301667 | 55629561 | 837400 | 20674000 | 160706 | 72.0 | 1.1 | 26.7 | 0.2 |
| | reBE | 73955727 | 53798759 | 931117 | 19061468 | 164383 | 72.7 | 1.3 | 25.8 | 0.2 |
| 8 | CD | 94242000 | 69981194 | 895086 | 21924879 | 1440841 | 74.3 | 0.9 | 23.3 | 1.5 |
| | CE | 87521433 | 65826096 | 795174 | 19898493 | 1001670 | 75.2 | 0.9 | 22.7 | 1.1 |

45 samples

| | | | | | | | | | |
|---|---|---|---|---|---|---|---|---|---|
| CF | 78577123 | 57350715 | 727499 | 19370047 | 1128862 | 73.0 | 0.9 | 24.7 | 1.4 |
| CG | 74335662 | 54031409 | 757165 | 18219477 | 1327611 | 72.7 | 1.0 | 24.5 | 1.8 |
| CH | 71311090 | 51495180 | 600311 | 18212945 | 1002654 | 72.2 | 0.8 | 25.5 | 1.4 |
| CI | 68127601 | 50892326 | 676629 | 15685180 | 873466 | 74.7 | 1.0 | 23.0 | 1.3 |
| CJ | 76715038 | 56976530 | 709022 | 17405444 | 1624042 | 74.3 | 0.9 | 22.7 | 2.1 |
| CK | 86413278 | 61304595 | 792195 | 21882879 | 2433609 | 70.9 | 0.9 | 25.3 | 2.8 |
| O | 65291420 | 48553707 | 643984 | 15921217 | 172512 | 74.4 | 1.0 | 24.4 | 0.3 |
| AX | 81707619 | 54047977 | 806730 | 26666168 | 186744 | 66.1 | 1.0 | 32.6 | 0.2 |
| BA | 76840312 | 54216509 | 717471 | 21683193 | 223139 | 70.6 | 0.9 | 28.2 | 0.3 |
| M | 68203557 | 52354773 | 705256 | 14871585 | 271943 | 76.8 | 1.0 | 21.8 | 0.4 |
| AJ | 74415306 | 54943719 | 853306 | 18434630 | 183651 | 73.8 | 1.1 | 24.8 | 0.2 |
| BN | 71820126 | 52418700 | 705993 | 18417595 | 277838 | 73.0 | 1.0 | 25.6 | 0.4 |
| BO | 77999787 | 54203706 | 801690 | 22772556 | 221835 | 69.5 | 1.0 | 29.2 | 0.3 |
| L | 73132084 | 53677667 | 863053 | 18353070 | 238294 | 73.4 | 1.2 | 25.1 | 0.3 |
| U | 73354711 | 52946806 | 735464 | 19522634 | 149807 | 72.2 | 1.0 | 26.6 | 0.2 |
| AL | 65449989 | 49489112 | 706504 | 15025702 | 228671 | 75.6 | 1.1 | 23.0 | 0.3 |
| AA | 74004023 | 54406152 | 728663 | 18666010 | 203198 | 73.5 | 1.0 | 25.2 | 0.3 |
| Z | 82069429 | 42530992 | 757955 | 38615610 | 164872 | 51.8 | 0.9 | 47.1 | 0.2 |
| AM | 74445255 | 53816118 | 708349 | 19751860 | 168928 | 72.3 | 1.0 | 26.5 | 0.2 |
| BF | 75293665 | 55313244 | 729147 | 19021215 | 230059 | 73.5 | 1.0 | 25.3 | 0.3 |
| BJ | 75860791 | 55080260 | 796528 | 19810316 | 173687 | 72.6 | 1.0 | 26.1 | 0.2 |
| BK | 75078043 | 55261371 | 728620 | 18925556 | 162496 | 73.6 | 1.0 | 25.2 | 0.2 |
| N | 73051151 | 54625170 | 788424 | 17440856 | 196701 | 74.8 | 1.1 | 23.9 | 0.3 |
| P | 63910979 | 47182350 | 773856 | 15772964 | 181809 | 73.8 | 1.2 | 24.7 | 0.3 |
| BD | 73821468 | 55838692 | 835082 | 16931934 | 215760 | 75.6 | 1.1 | 22.9 | 0.3 |
| BE | 63107622 | 46862674 | 792590 | 15285084 | 167274 | 74.3 | 1.3 | 24.2 | 0.3 |
| R | 74926891 | 55205551 | 773100 | 18764622 | 183618 | 73.7 | 1.0 | 25.0 | 0.2 |
| AI | 82678733 | 52219876 | 693404 | 29629619 | 135834 | 63.2 | 0.8 | 35.8 | 0.2 |
| BV | 75559608 | 52931605 | 665076 | 21792244 | 170683 | 70.1 | 0.9 | 28.8 | 0.2 |
| A | 76404110 | 53049371 | 718347 | 22486438 | 149954 | 69.4 | 0.9 | 29.4 | 0.2 |
| S | 74955482 | 53813012 | 756326 | 20228332 | 157812 | 71.8 | 1.0 | 27.0 | 0.2 |
| CC | 64623480 | 47161637 | 653520 | 16657057 | 151266 | 73.0 | 1.0 | 25.8 | 0.2 |
| Y | 90297268 | 41308218 | 617902 | 48221568 | 149580 | 45.7 | 0.7 | 53.4 | 0.2 |
| AC | 76488390 | 53781098 | 702343 | 21772489 | 232460 | 70.3 | 0.9 | 28.5 | 0.3 |
| BI | 73934733 | 53636277 | 671656 | 19484276 | 142524 | 72.5 | 0.9 | 26.4 | 0.2 |
| AO | 74617649 | 55997702 | 664336 | 17764577 | 191034 | 75.0 | 0.9 | 23.8 | 0.3 |
| BQ | 73547358 | 55322600 | 582259 | 17438305 | 204194 | 75.2 | 0.8 | 23.7 | 0.3 |
| BT | 67672343 | 50043531 | 514074 | 16969694 | 145044 | 73.9 | 0.8 | 25.1 | 0.2 |
| BC | 74022020 | 55179346 | 723468 | 17873546 | 245660 | 74.5 | 1.0 | 24.1 | 0.3 |
| BX | 72391959 | 53279904 | 958694 | 17960936 | 192425 | 73.6 | 1.3 | 24.8 | 0.3 |
| BY | 71979863 | 53911154 | 685654 | 17189959 | 193096 | 74.9 | 1.0 | 23.9 | 0.3 |
| W | 73160618 | 52485318 | 870087 | 19638185 | 167028 | 71.7 | 1.2 | 26.8 | 0.2 |
| X | 74463403 | 52400691 | 898214 | 20959981 | 204517 | 70.4 | 1.2 | 28.1 | 0.3 |
| BW | 73307409 | 52745754 | 837396 | 19589668 | 134591 | 72.0 | 1.1 | 26.7 | 0.2 |
| CA | 73516158 | 52906856 | 846636 | 19583046 | 179620 | 72.0 | 1.2 | 26.6 | 0.2 |
| AE | 218114881 | 17716958 | 1196469 | 199149566 | 51888 | 8.1 | 0.5 | 91.3 | 0.0 |
| AF | 72108676 | 53473615 | 715719 | 17735243 | 184099 | 74.2 | 1.0 | 24.6 | 0.3 |
| AR | 71885923 | 53443413 | 725863 | 17540971 | 175676 | 74.3 | 1.0 | 24.4 | 0.2 |
| AH | 220438330 | 16951685 | 1160537 | 202281283 | 44825 | 7.7 | 0.5 | 91.8 | 0.0 |
| AS | 71477049 | 53095094 | 774339 | 17408436 | 199180 | 74.3 | 1.1 | 24.4 | 0.3 |
| AT | 71468031 | 53591317 | 770686 | 16885869 | 220159 | 75.0 | 1.1 | 23.6 | 0.3 |
| | | | | | mean(-outliers) | 73.2 | 1.0 | 25.3 | 0.5 |
| | | | | | excluding A & BO: | 73.3 | 1.0 | 25.2 | 0.5 |

*Samples are colored by library preparation batch. Uncolored samples were excluded. The columns for “total reads” and “total assigned” are conditionally colored to highlight, in the first one, lower (green) and higher (red) values; *vice-versa* in the second one. Criteria for inclusion were “> 70% assigned from total reads” (9 samples highlighted in gray, from which samples A and BO were accepted), and/or “< 2SD % multi-mapped” (5 samples highlighted in gray). Therefore, 7 samples were excluded. Samples AF, BX, BE, BK, were replaced by newly sequenced samples reAF, reBX, reBE, reBK, to compensate for batch effects.

**Table S5.** List of ribosomal genes excluded from the gene count matrix*.

*RN7SK, RN7SKP10, RN7SKP104, RN7SKP109, RN7SKP11, RN7SKP122, RN7SKP129, RN7SKP15, RN7SKP16, RN7SKP160, RN7SKP163, RN7SKP17, RN7SKP170, RN7SKP172, RN7SKP173, RN7SKP176, RN7SKP177, RN7SKP188, RN7SKP197, RN7SKP203, RN7SKP208, RN7SKP214, RN7SKP225, RN7SKP229, RN7SKP23, RN7SKP237, RN7SKP241, RN7SKP243, RN7SKP250, RN7SKP26, RN7SKP268, RN7SKP269, RN7SKP271, RN7SKP272, RN7SKP275, RN7SKP287, RN7SKP288, RN7SKP30, RN7SKP34, RN7SKP36, RN7SKP45, RN7SKP49, RN7SKP66, RN7SKP70, RN7SKP71, RN7SKP74, RN7SKP78, RN7SKP80, RN7SKP87, RN7SKP9, RN7SKP97, RN7SL1, RN7SL113P, RN7SL125P, RN7SL127P, RN7SL130P, RN7SL134P, RN7SL145P, RN7SL152P, RN7SL15P, RN7SL167P, RN7SL172P, RN7SL181P, RN7SL187P, RN7SL192P, RN7SL2, RN7SL205P, RN7SL208P, RN7SL209P, RN7SL20P, RN7SL213P, RN7SL216P, RN7SL217P, RN7SL225P, RN7SL229P, RN7SL234P, RN7SL236P, RN7SL239P, RN7SL23P, RN7SL244P, RN7SL253P, RN7SL25P, RN7SL263P, RN7SL270P, RN7SL272P, RN7SL273P, RN7SL275P, RN7SL279P, RN7SL296P, RN7SL3, RN7SL300P, RN7SL306P, RN7SL320P, RN7SL333P, RN7SL336P, RN7SL338P, RN7SL341P, RN7SL34P, RN7SL351P, RN7SL357P, RN7SL35P, RN7SL361P, RN7SL364P, RN7SL368P, RN7SL369P, RN7SL370P, RN7SL377P, RN7SL381P, RN7SL388P, RN7SL391P, RN7SL395P, RN7SL398P, RN7SL403P, RN7SL40P, RN7SL42P, RN7SL430P, RN7SL431P, RN7SL434P, RN7SL43P, RN7SL443P, RN7SL448P, RN7SL470P, RN7SL477P, RN7SL479P, RN7SL481P, RN7SL484P, RN7SL487P, RN7SL49P, RN7SL4P, RN7SL510P, RN7SL517P, RN7SL521P, RN7SL526P, RN7SL535P, RN7SL559P, RN7SL566P, RN7SL573P, RN7SL574P, RN7SL577P, RN7SL589P, RN7SL591P, RN7SL592P, RN7SL5P, RN7SL605P, RN7SL608P, RN7SL614P, RN7SL634P, RN7SL648P, RN7SL650P, RN7SL652P, RN7SL657P, RN7SL65P, RN7SL663P, RN7SL66P, RN7SL671P, RN7SL674P, RN7SL677P, RN7SL684P, RN7SL689P, RN7SL68P, RN7SL6P, RN7SL704P, RN7SL717P, RN7SL724P, RN7SL731P, RN7SL732P, RN7SL736P, RN7SL737P, RN7SL738P, RN7SL749P, RN7SL751P, RN7SL752P, RN7SL75P, RN7SL762P, RN7SL767P, RN7SL76P, RN7SL784P, RN7SL789P, RN7SL793P, RN7SL800P, RN7SL806P, RN7SL809P, RN7SL812P, RN7SL81P, RN7SL820P, RN7SL825P, RN7SL826P, RN7SL827P, RN7SL832P, RN7SL834P, RN7SL840P, RN7SL850P, RN7SL870P, RN7SL8P, RN7SL92P, RNA18SN1, RNA18SN1_1, RNA18SN2, RNA18SN3, RNA18SN3_1, RNA18SN4, RNA18SN5, RNA18SP, RNA18SP3, RNA18SP4, RNA18SP5, RNA28SN1, RNA28SN1_1, RNA28SN2, RNA28SN3, RNA28SN3_1, RNA28SN4, RNA28SN5, RNA28SP, RNA45SN1, RNA45SN1_1, RNA45SN2, RNA45SN3, RNA45SN3_1, RNA45SN4, RNA45SN5, RNA5-8SN1, RNA5-8SN1_1, RNA5-8SN2, RNA5-8SN3, RNA5-8SN4, RNA5-8SP3, RNA5-8SP4, RNA5-8SP8, RNA5S9, RNA5SP108, RNA5SP118, RNA5SP122, RNA5SP123, RNA5SP132, RNA5SP150, RNA5SP151, RNA5SP152, RNA5SP155, RNA5SP159, RNA5SP160, RNA5SP162, RNA5SP18, RNA5SP184, RNA5SP187, RNA5SP202, RNA5SP203, RNA5SP21, RNA5SP216, RNA5SP217, RNA5SP219, RNA5SP236, RNA5SP244, RNA5SP265, RNA5SP280, RNA5SP283, RNA5SP296, RNA5SP317, RNA5SP322, RNA5SP323, RNA5SP325, RNA5SP352, RNA5SP37, RNA5SP379, RNA5SP387, RNA5SP392, RNA5SP393, RNA5SP422, RNA5SP424, RNA5SP434, RNA5SP462, RNA5SP464, RNA5SP473, RNA5SP474, RNA5SP477, RNA5SP490, RNA5SP493, RNA5SP505, RNA5SP530, RNA5SP82, RNA5SP91, RNA5SP92*

*Only genes with sum counts ≥ 10 across all samples are shown in the list.

**Table S6.** Poly-acrylamide proportions in hydrogels at different stiffnesses used for the migration assay.

| Stiffness | Acrylamide % | Bis-Acrylamide % | Reference |
|---|---|---|---|
| 500 Pa | 8 | 0.01 | [4] |
| 3.2 kPa | 4 | 0.3 | [1] |
| 4.5 kPa | 5 | 0.15 | [1] |
| 10 kPa | 7.5 | 0.35 | [5] |
| 35 kPa | 10 | 0.2 | [6] |

**Table S7.** Real Time quantitative PCR primer sequences.

| Gene | Forward primer | Reverse primer |
|---|---|---|
| *YAP1* | *ACG ATG CCC TCT GTA CTG AC* | *ACA AGA GAC CAC ATC AAG GC* |
| *MGLL* | *CTG GTG GGT GCT CTG AAA AC* | *GAA GCC ATC GTG ATC CCA AC* |
| *MATN3* | *ACT GAG GAA GCA CGA AGA CT* | *GAG CTG ACC TTG TCC TGG AA* |
| *KCTD20* | *ATT TCT ATG CCC CTC CCA CC* | *CTC AAC CCA GCC CTT GTT TC* |
| *CLEC3B* | *GCA TCG CCT ACA AGA ACT GG* | *CTT GTC GAA CCA CTT GCC G* |
| *GAB1* | *AGT CAG CAC TTT GGT CCA CT* | *AGA TAG CCT CAC CCT ACC CA* |

**Table S8.** Supporting literature on process involvement of genes categorized in specific LRT subsets.

| Gene Name | Categorization | Cellular Process | Reference |
|---|---|---|---|
| *SESN3* | Unique to SS | Cellular antioxidant processes | Kim H, An S, Ro S-H, Teixeira F, Park GJ, Kim C, et al. Janus-faced Sestrin2 controls ROS and mTOR signalling through two separate functional domains. *Nat Commun* 2015;6. |
| *TALDO1* | Unique to SS | Cellular antioxidant processes | Perl A, Hanczko R, Telarico T, Oaks Z, Landas S. Oxidative stress, inflammation and carcinogenesis are controlled through the pentose phosphate pathway by transaldolase. *Trends Mol Med* 2011;17:395–403. |
| *NQO1* | Unique to SS | Cellular antioxidant processes | Nishida-Tamehiro K, Kimura A, Tsubata T, Takahashi S, Suzuki Id H. Antioxidative enzyme NAD(P)H quinone oxidoreductase 1 (NQO1) modulates the differentiation of Th17 cells by regulating ROS levels. *PLoS One* 2022;17:e0272090. |
| *TGM2* | Unique to SS | ECM modulation | Faye C, Inforzato A, Bignon M, Hartmann DJ, Muller L, Ballut L, et al. Transglutaminase-2: a new endostatin partner in the extracellular matrix of endothelial cells. *Biochem J* 2010;427:467. |
| *ADAMTS18* | Unique to SS | ECM modulation | Ataca D, Aouad P, Constantin C, Laszlo C, Beleut M, Shamseddin M, et al. The secreted protease Adamts18 links hormone action to activation of the mammary stem cell niche. *Nat Commun* 2020;11. |
| *CLDN10* | Unique to SS | Cell Adhesion | Tsukita S, Furuse M. The structure and function of claudins, cell adhesion molecules at tight junctions. *Ann N Y Acad Sci* 2000;915:129–135. |
| *CLDN11* | Unique to SS | Cell Adhesion | Tsukita S, Furuse M. The structure and function of claudins, cell adhesion molecules at tight junctions. *Ann N Y Acad Sci* 2000;915:129–135. |
| *NR2F1* | Unique to SS | Transcription factor | Not available |
| *FLII* | Unique to SFN | Cytoskeleton reorganization | Mohammad I, Arora PD, Naghibzadeh Y, Wang Y, Li J, Mascarenhas W, et al. Flightless I is a focal adhesion-associated actin-capping protein that regulates cell migration. *The FASEB Journal* 2012;26:3260–3272. |
| *PPP1R16A* | Unique to SFN | Cytoskeleton reorganization | Grassie ME, Moffat LD, Walsh MP, Macdonald JA. The myosin phosphatase targeting protein (MYPT) family: A regulated mechanism for achieving substrate specificity of the catalytic subunit of protein phosphatase type 1d. *Arch Biochem Biophys* 2011;510:147–159. |
| *RABGAP1L* | Unique to SFN | Vesicle transport | Qu F, Lorenzo DN, King SJ, Brooks R, Bear JE, Bennett V. Ankyrin-B is a PI3P effector that promotes polarized a5b1-integrin recycling via recruiting RabGAP1L to early endosomes. *Elife* 2016;5:e20417. |
| *CDK16* | Unique to SFN | Vesicle transport | Dohmen M, Krieg S, Agalaridis G, Zhu X, Shehata SN, Pfeiffenberger E, et al. AMPK-dependent activation of the Cyclin Y/CDK16 complex controls autophagy. *Nat Commun* 2020;11. |
| *MAP2K3* | Unique to SFN | Kinase | Not available |
| *TYK2* | Unique to SFN | Kinase | Not available |
| *GADD45B* | Unique to SFN | Kinase | Not available |
| *ATP1B1* | Present in SS, SFN and INT | Cytoskeleton modulation | Bab-Dinitz E, Albeck S, Peleg Y, Brumfeld V, Gottschalk KE, Karlish SJD. A C-terminal lobe of the β subunit of Na,K- |

| | | | |
|---|---|---|---|
| | | | ATPase and H,K-ATPase resembles cell adhesion molecules. *Biochemistry* 2009;48:8684–8691. |
| *TAL1* | Present in SS, SFN and INT | Cytoskeleton modulation | Obeng G, Park EJ, Appiah MG, Kawamoto E, Gaowa A, Shimaoka M. miRNA-200c-3p targets talin-1 to regulate integrin-mediated cell adhesion. *Scientific Reports* \| 123AD;11:21597. |
| *SYNGR2* | Present in SS, SFN and INT | Cytoskeleton modulation | Francis MS, Lai C-H, Mirey G, Jank T, Shenker BJ, H-y T, et al. Internalization of the Active Subunit of the Aggregatibacter actinomycetemcomitans Cytolethal Distending Toxin Is Dependent upon Cellugyrin (Synaptogyrin 2), a Host Cell Non-Neuronal Paralog of the Synaptic Vesicle Protein, Synaptogyrin 1. *Front Cell Infect Microbiol* 2017;7:469. |
| *PICK1* | Present in SS, SFN and INT | Cytoskeleton modulation | Ramsakha N, Ojha P, Pal S, Routh S, Citri A, Bhattacharyya S. A vital role for PICK1 in the differential regulation of metabotropic glutamate receptor internalization and synaptic AMPA receptor endocytosis. *J Biol Chem* 2023;299:104837. |
| *TRIM46* | Present in SS, SFN and INT | Cytoskeleton modulation | Harterink M, Vocking K, Pan X, Soriano Jerez EM, Slenders L, Fréal A, et al. TRIM46 Organizes Microtubule Fasciculation in the Axon Initial Segment. *J Neurosci* 2019;39:4864–4873. |
| *CXCL8* | Present in SS, SFN and INT | Inflammation | Not available |
| *IL6* | Present in SS, SFN and INT | Inflammation | Not available |
| *MAP4K4* | Present in SS, SFN and INT | Inflammation | Not available |
| *PTX3* | Present in SS, SFN and INT | Inflammation | Zlibut A, Bocsan IC, Agoston-Coldea L. Pentraxin-3 and endothelial dysfunction. *Adv Clin Chem*. Academic Press Inc.; 2019. p163–179. |
| *TRAFD1* | Present in SS, SFN and INT | Inflammation | Sanada T, Takaesu G, Mashima R, Yoshida R, Kobayashi T, Yoshimura A. FLN29 Deficiency Reveals Its Negative Regulatory Role in the Toll-like Receptor (TLR) and Retinoic Acid-inducible Gene I (RIG-I)-like Helicase Signaling Pathway. *J Biol Chem* 2008;283:33858–33864. |
| *RNLS* | Present in SS, SFN and INT | Enzymes involved hydrolysis | Vijayakumar A, Mahapatra NR. Renalase: a novel regulator of cardiometabolic and renal diseases. *Hypertension Res* 2022;45:1582–1598. |
| *ABHD11* | Present in SS, SFN and INT | Enzymes involved hydrolysis | Arya M, Srinivasan M, Rajasekharan R. Human alpha beta hydrolase domain containing protein 11 and its yeast homolog are lipid hydrolases. *Biochem Biophys Res Commun* 2017;487:875–880. |
| *HOGA1* | Present in SS and INT, not in SFN | Mitochondrial respiration | Belostotsky R, Seboun E, Idelson GH, Milliner DS, Becker-Cohen R, Rinat C, et al. Mutations in DHDPSL Are Responsible For Primary Hyperoxaluria Type III. *Am J Hum Genet* 2010;87:392–399. |
| *PLSCR3_1* | Present in SS and INT, not in SFN | Mitochondrial respiration | Liu J, Epand RF, Durrant D, Grossman D, Chi NW, Epand RM, et al. Role of phospholipid scramblase 3 in the regulation of tumor necrosis factor-α-induced apoptosis. *Biochemistry* 2008;47:4518–4529. |
| *NDUFB10* | Present in SS and INT, not in SFN | Mitochondrial respiration | Arroum T, Borowski MT, Marx N, Schmelter F, Scholz M, Psathaki OE, et al. Loss of respiratory complex i subunit |

| | | | |
|---|---|---|---|
| | | | NDUFB10 affects complex i assembly and supercomplex formation. *Biol Chem* 2023;404:399–415. |
| *HIGD2A* | Present in SS and INT, not in SFN | Mitochondrial respiration | Hock DH, Reljic B, Ang CS, Muellner-Wong L, Mountford HS, Compton AG, et al. HIGD2A is Required for Assembly of the COX3 Module of Human Mitochondrial Complex IV. *Mol Cell Proteomics* 2020;19:1145. |
| *VPS13D* | Present in SS and INT, not in SFN | Mitochondrial respiration | Guillén-Samander A, Leonzino M, Hanna MG, Tang N, Shen H, Camilli P De. VPS13D bridges the ER to mitochondria and peroxisomes via Miro. *J Cell Biol* 2021;220. |
| *NDUFA9* | Present in SS and INT, not in SFN | Mitochondrial respiration | Stroud DA, Formosa LE, Wijeyeratne XW, Nguyen TN, Ryan MT. Gene Knockout Using Transcription Activator-like Effector Nucleases (TALENs) Reveals That Human NDUFA9 Protein Is Essential for Stabilizing the Junction between Membrane and Matrix Arms of Complex I. *J Biol Chem* 2013;288:1685. |
| *PCK2* | Present in SS and INT, not in SFN | Mitochondrial respiration | Bluemel G, Planque M, Madreiter-Sokolowski CT, Haitzmann T, Hrzenjak A, Graier WF, et al. PCK2 opposes mitochondrial respiration and maintains the redox balance in starved lung cancer cells. *Free Radic Biol Med* 2021;176:34–45. |
| *LSS_1* | Present in SS and INT, not in SFN | Cholesterol biosynthesis or transport | Thoma R, Schulz-Gasen T, D'Arcy B, Benz J, Aebi J, Dehmlow H, et al. Insight into steroid scaffold formation from the structure of human oxidosqualene cyclase. *Nature* 2004 432:7013 2004;432:118–122. |
| *OSBPL6* | Present in SS and INT, not in SFN | Cholesterol biosynthesis or transport | Ouimet M, Hennessy EJ, Solingen C Van, Koelwyn GJ, Hussein MA, Ramkhelawon B, et al. miRNA Targeting of Oxysterol Binding Protein-like 6 (OSBPL6) Regulates Cholesterol Trafficking and Efflux. *Arterioscler Thromb Vasc Biol* 2016;36:942. |
| *EIF5A* | Present in SS and INT, not in SFN | Cytoskeleton dynamics | Muñoz-Soriano V, Domingo-Muelas A, Li T, Gamero E, Bizy A, Fariñas I, et al. Evolutionary conserved role of eukaryotic translation factor eIF5A in the regulation of actin-nucleating formins. *Sci Rep* 2017;7. |
| *TUBB_3* | Present in SS and INT, not in SFN | Cytoskeleton dynamics | Isrie M, Breuss M, Tian G, Hansen AH, Cristofoli F, Morandell J, et al. Mutations in Either TUBB or MAPRE2 Cause Circumferential Skin Creases Kunze Type. *Am J Hum Genet* 2015;97:790. |
| *SLC44A3* | Present in SS and INT, not in SFN | Transmembrane transport | Traiffort E, O'regan S, Ruat M. The choline transporter-like family SLC44: Properties and roles in human diseases q. *Mol Aspects Med* 2013;34:646–654. |
| *CD79B* | Present in SS and INT, not in SFN | Immune response | Huse K, Bai B, Hilden VI, Bollum LK, Våtsveen TK, Munthe LA, et al. Mechanism of CD79A and CD79B Support for IgM+ B Cell Fitness through B Cell Receptor Surface Expression. *J Immunol* 2022;209:2042–2053. |
| *SELE* | Present in SS and INT, not in SFN | Immune response | McEver RP. Selectins: initiators of leucocyte adhesion and signalling at the vascular wall. *Cardiovasc Res* 2015;107:331. |
| *IKZF1* | Present in SS and INT, not in SFN | Transcription factor | Not available |

| | | | |
|---|---|---|---|
| *ZBTB3* | Present in SS and INT, not in SFN | Transcription factor | Not available |
| *CALR* | Both SS and SFN, but no INT | Protein folding | Venkatesan A, Satin LS, Raghavan M. Roles of Calreticulin in Protein Folding, Immunity, Calcium Signaling and Cell Transformation. *Prog Mol Subcell Biol* 2021;59:145–162. |
| *CANX* | Both SS and SFN, but no INT | Protein folding | Wang H, Li S, Wang J, Chen S, Sun XL, Wu Q. N-glycosylation in the protease domain of trypsin-like serine proteases mediates calnexin-assisted protein folding. *Elife* 2018;7. |
| *AFG1L* | Both SS and SFN, but no INT | Mitochondrial enzyme | Germany EM, Zahayko N, Huebsch ML, Fox JL, Prahlad V, Khalimonchuk O. The AAA ATPase Afg1 preserves mitochondrial fidelity and cellular health by maintaining mitochondrial matrix proteostasis. *J Cell Sci* 2018;131. |
| *THEM5* | Both SS and SFN, but no INT | Mitochondrial enzyme | Zhuravleva E, Gut H, Hynx D, Marcellin D, Bleck CKE, Genoud C, et al. Acyl Coenzyme A Thioesterase Them5/Acot15 Is Involved in Cardiolipin Remodeling and Fatty Liver Development. *Mol Cell Biol* 2012;32:2685. |
| *PINK1* | Both SS and SFN, but no INT | Mitochondrial enzyme | Yang Y, Ouyang Y, Yang L, Beal MF, McQuibban A, Vogel H, et al. Pink1 regulates mitochondrial dynamics through interaction with the fission/fusion machinery. *Proc Natl Acad Sci U S A* 2008;105:7070–7075. |
| *CDK9* | Both SS and SFN, but no INT | Transcription regulatory enzymes | Brauns-Schubert P, Schubert F, Wissler M, Weiss M, Schlicher L, Bessler S, et al. CDK9-mediated phosphorylation controls the interaction of TIP60 with the transcriptional machinery. *EMBO Rep* 2018;19:244–256. |
| *GPX3* | Both SS and SFN, but no INT | Antioxidant enzymes | Li Y, Zhou Y, Liu D, Wang Z, Qiu J, Zhang J, et al. Glutathione Peroxidase 3 induced mitochondria-mediated apoptosis via AMPK /ERK1/2 pathway and resisted autophagy-related ferroptosis via AMPK/mTOR pathway in hyperplastic prostate. *J Transl Med* 2023;21. |
| *CCL2* | Both SS and SFN, but no INT | Chemotactic cytokines | Not available |

**Table S9.** Top processes obtained by Ingenuity Pathway Analysis (IPA) from the 603 interaction genes.

| | *p*-value ranges |
|---|---|
| **Top molecular and cellular functions** | |
| Cell death and survival | 4.74E-04 – 2.05E-08 |
| Cellular movement | 4.18E-04 – 1.01E-07 |
| Cellular response to therapeutics | 5.24E-05 – 3.05E-07 |
| Carbohydrate metabolism | 1.11E-06 – 1.11E-06 |
| **Top physiological system development and function processes** | |
| Embryonic development | 4.75E-04 – 1.01E-07 |
| Cardiovascular system development and function | 4.42E-04 – 3.80E-06 |
| Tissue development | 4.42E-04 – 3.80E-06 |

**Table S10.** Genes supporting the IPA results for migration at 5 dynes/cm$^2$ per SFN. IPA activation predictions considered significant when $|z| \geq 2$.

| kPa | Z-score (IPA) | Predicted | # genes migration EC | list |
|---|---|---|---|---|
| 100 | -2.613 | Decreased | 78 | *ACKR3,ADAM15,ADGRA2,ANGPTL4,ARRB2,BCAR1,BMX,C5,CAV1,CAVIN2,CCL2,CD63,CD81,CRYAB,CXCL1,CYTOR,DPP4,ELN,EPHB4,ETS1,EXOC3L2,F11R,F3,FGF2,FLT1,FOXC1,FOXC2,FOXO1,FOXP1,GATA2,GATA3,GATA6,HGF,HLX,ID1,IGF2,IL33,ITGB1BP1,ITGB3,ITGB4,KLF2,LIPA,MAP2K1,MAP2K6,MAPKAPK5,MET,NOX4,NOXA1,NR4A1,PDGFB,PGF,PIK3C2B,PIK3CG,PLD1,PPARG,PRKCE,PROX1,RAB7A,RGCC,S1PR3,SDC4,SERPIND1,SERPINE1,SH2D2A,SMAD3,SP1,SPRY4,STAB2,TEK,TFPI,THSD7A,TNFRSF6B,TNFSF15,TNS3,VAV3,VCAM1,VEGFC,YAP1* |
| 10 | -1.688 | · | 31 | *ACKR3,BCAR1,BMX,CAVIN2,CD63,CD81,CRYAB,CYTOR,EXOC3L2,F3,FLT1,FOXO1,GATA2,GATA6,HGF,HLX,IL33,ITGB1BP1,ITGB4,MET,NOX4,PGF,RGCC,RNF7,S1PR3,SERPIND1,SMAD3,SPRY4,TGFB3,THSD7A,TNS3* |
| 1 | -3.212 | Decreased | 122 | *ACKR3,ACP1,ADAM15,ADGRA2,AMOT,ANGPT1,ANGPTL4,ANXA3,ARFGEF2,ARHGEF6,ARRB2,ATP2B4,BCAR1,BDKRB2,BDNF,BMP4,BMX,BTC,C5,CAV1,CAVIN2,CCL2,CD36,CD63,CD81,CD82,CDH13,CDKN1B,CRYAB,CXCL1,CXCL12,CYTOR,DICER1,DIP2A,DPP4,EPHB4,F11R,F3,FGF2,FLT1,FOXC1,FOXC2,FOXO1,FOXO3,FOXP1,G6PD,GATA2,GATA3,GATA6,HGF,HLX,HMMR,HRAS,HSPB1,IGFBP3,IL33,ITGB1BP1,ITGB3,ITGB4,KLF2,KMT2A,LGALS1,LIPA,MAP2K1,MAP2K6,MAPKAPK5,MEOX2,MET,NFATC2,NFATC3,NOS3,NOX4,NOXA1,NR4A1,NRG1,NRP1,PDGFB,PDGFC,PGF,PIK3C2B,PIK3CG,PLD1,PRKCE,PROS1,PROX1,PTK2,RAB7A,RAC1,RECK,RGCC,RNF7,ROBO1,ROBO2,S100A2,S1PR3,SASH1,SDC4,SERPIND1,SERPINE1,SH2D2A,SLIT2,SMAD3,SP1,SPRY4,STAB2,TEK,TFPI,TGFB2,TGFB3,TGFBI,THRA,THSD7A,TIMP3,TJP1,TMSB10/TMSB4X,TNFRSF12A,TNFRSF6B,TNFSF15,TNS3,VCAM1,VEGFC,YAP1* |

**Table S11.** Example genes known to be modulated by YAP1 (ChEA database)[7] and which are present in the LRT interaction list. Indicated are their adj-*p* (FDR).

| target | FDR (Int) | target | FDR (Int) | target | FDR (Int) | target | FDR (Int) | target | FDR (Int) |
|---|---|---|---|---|---|---|---|---|---|
| *TACC1* | 0.0022 | *KRT15* | 0.0145 | *PEAK1* | 0.0232 | *STIM2* | 0.027 | *TAC4* | 0.0330 |
| *SYNGR2* | 0.0023 | *CCN1* | 0.016 | *LRBA* | 0.0242 | *BAIAP2L1* | 0.0274 | *RHOB* | 0.0337 |
| *TRAFD1* | 0.0037 | *RBM33* | 0.0168 | *AGO2* | 0.0252 | *ADORA2B* | 0.0286 | *KMT2C* | 0.0341 |
| *PTX3* | 0.0038 | *FOXF1* | 0.019 | *EML4* | 0.0252 | *STRADB* | 0.0288 | *GBX2* | 0.0366 |
| *DTNA* | 0.0084 | *RNF32* | 0.0203 | *YWHAE* | 0.0252 | *ASNS* | 0.0309 | *PPM1L* | 0.0370 |
| *MME* | 0.0084 | *SEMA6A* | 0.0204 | *DOCK10* | 0.026 | *HSPA5* | 0.0317 | *MAMDC2* | 0.0388 |
| *TRIM44* | 0.0084 | *HNRNPL_1* | 0.0212 | *USP42* | 0.0262 | *CDYL2* | 0.0323 | *RAPGEF2* | 0.0400 |
| *DOCK4* | 0.011 | *TUBB3* | 0.0214 | *TPD52* | 0.0268 | *KDM5B* | 0.0323 | *GAB1* | 0.0408 |

### Supplementary Files (available after publication in journal)

#### Supplementary Movie 1 (separate file)

Animated IPA network highlighting changes in gene regulation and predicted process activation across the 11 non-static conditions.

#### Supplementary Data 1 (separate file)

DESeq2 results of the LRT tests for shear stress, stiffness and the interaction term in the full equation vs. respective reduced equations. Tables include the complete gene lists with adjusted *p*-values. Unique gene lists after excluding overlapped genes between groups are included.

#### Supplementary Data 2 (separate file)

DESeq2 results of the Wald test comparing all conditions vs. the stiffest (100 kPa) static condition as the reference. Tables include the complete gene lists with fold changes and adjusted *p*-values.

#### Supplementary Data 3 (separate file)

DESeq2 results of the Wald test comparing all conditions to their own static condition per each stiffness group as the reference. Tables include the complete gene lists with fold changes and adjusted *p*-values.

#### Supplementary Data 4 (separate file)

Worksheet 1: KEGG pathways ranked by differential activation between substrate stiffness conditions (ΔSFN) at 25 and at 40 dynes/$cm^2$ shear stress. ΔSFN was calculated as mean $log_2$FC (100 kPa) – mean $log_2$FC (10 kPa). The number of contributing genes is indicated for each pathway. Pathways displayed in Figure 4B are highlighted in gray.

Worksheet 2: Genes contributing to the divergence between angiogenic profiles across mechanical conditions, ranked by ΔMech = $log_2$FC (40 dynes/$cm^2$, 10 kPa) − $log_2$FC (25 dynes/$cm^2$, 100 kPa). Analysis was restricted to genes belonging to the top KEGG pathway groups identified in the pathway-level analysis (see Supplementary Methods). Genes displayed in Figure 4C are highlighted in gray.

#### Supplementary Data 5 (separate file)

Worksheet 1: List of the 603 INT-genes and their Pearson correlation values of gene expression with active YAP1 nuclear intensity (N/C). The 5 genes selected for further exploration are highlighted in gray.

Worksheet 2: *YAP1* knockdown ΔCt values for the selected candidate genes across combined SS and substrate SFN conditions.

Worksheet 3: *YAP1* knockdown effect size (ΔΔCt) and confidence interval values for the selected genes across combined SS and substrate SFN conditions.

### Supplementary Methods

#### *Hydrogel preparation at different SFNs*

Glass slides were treated with 20% (3-Aminopropyl)trimethoxysilane (APTS; 281778, Sigma-Aldrich) for 5 min, washed twice in abundant sterile deionized water, and allowed to dry. Slides were then treated with 1% glutaraldehyde (G6257, Sigma-Aldrich) for 30 min, washed twice in abundant sterile deionized water, and again dried. In parallel, 1, 10 and 100 kPa hydrogel solutions were prepared by mixing acrylamide and bis-acrylamide (1610140 and 1610142, BioRad) in PBS at the percentages indicated in Table S1. Solutions were degassed inside the fume hood for 30 min and filtered through 0.2 µm filters.

For polymerization on slides, a BioRad gel casting system was adapted to hold three slides in between front and rear glasses. Four 3D-printed spacers were designed and printed in PETG. During functionalization and cell culture, hydrogels were subjected to incubations in DPBS and culture media. Polyacrylamide gels swell due to absorption of water, reaching a stable height after an overnight incubation. Different proportions of crosslinker may entail different swelling capacities. Therefore, a battery of hydrogel height measurements was performed to determine the height increase for each stiffness (1, 10, 100 kPa) before and after overnight incubations in DPBS. Accordingly, the 3D-printed spacers were designed for each gel stiffness such that after overnight incubations, hydrogel swelling resulted in exactly 2 mm height (Figure S1). The frontal glass of the casting system was treated with Sigmacote® (SL2, Sigma-Aldrich), a siliconizing reagent, to reduce its adherence to the hydrogel after polymerization.

Once the casting setup was mounted, TEMED and a 10% ammonium persulfate (APS) solution (both previously filtered through 0.2 µm filters) were added to the gel solutions at 1/1,000 and 1/100, respectively. These were gently resuspended and immediately pipetted into the gel casting system containing the slides, and hydrogels were let to polymerize for 40 min at RT. The hydrogels, still inside the gel casting glasses, were disassembled from the casting supports and placed in 150 mm Petri dishes with sterile Dulbecco's PBS (DPBS; Biowest). These were placed at 4°C overnight before functionalization.

#### *Hydrogel functionalization and cell seeding*

In the laminar hood, slides with hydrogels were disassembled from the casting glasses. Hydrogels were washed twice with abundant sterile DPBS. These were placed in CELLSTAR® FourWell plates (Greiner Bio-One) and treated with 0.5 mg/mL (in DPBS) Sulfo-SANPAH (sulfosuccinimidyl 6-(4′-azido-2′-nitrophenylamino)hexanoate); 803332, Sigma-Aldrich) for 20 min, and exposed to a 302 nm UV irradiation source that was placed 6 cm from the hydrogels. The solution was refreshed once, and the irradiation repeated. Hydrogels were gently washed twice in abundant sterile DPBS, and treated for 2h at 37°C with an 80 µg/mL solution of Collagen I from rat tail (Sigma-Aldrich) diluted in sterile 20 mM Acetic Acid. Functionalized hydrogels were gently washed twice in sterile DPBS and were incubated for 30 min in EGM-2MV containing 1x Antibiotic-Antimycotic (commercial mixture of penicillin, streptomycin and Amphotericin B, Gibco™) (from now on "media/AA").

HUVECs (C-12208, PromoCell) were used at passage P5-6 and grown in EC Growth Medium MV 2 (EGM-2MV; C-22121, PromoCell). HUVECs were seeded on the hydrogels at 0.25 M cells/hydrogel (8 hydrogels per each T75 flask), with 0.6 mL per hydrogel, carefully distributing the volume over the whole hydrogel surface. Cells were left to attach for 45 min to 1h at 37°C/5% $CO_2$, after which 4 mL media/AA was gently added to each well. HUVECs on hydrogels were incubated for at least 24h before any shear experiment.

*Shear stress experiments*

Hydrogels containing confluent HUVEC monolayers were placed in the bottom side of the custom flow chambers. A sterile silicon rubber gasket of 0.3 mm was added, surrounding the slide, and the top side of the chamber was mounted (Figure 1A). The flow-chamber system was placed at 37°C/5% $CO_2$ and was connected for 24h to a steady laminar flow of media using a peristaltic pump (Masterflex for 5 dynes/cm$^2$, NE-9000 for all other levels of shear stress). The pump programs included a ramping-up stage, starting from 10 dynes/cm$^2$ (35 mL/min) and gradually increasing flow rates to reach the desired SS level (Table S2). For static conditions (0 dynes/cm$^2$), hydrogels were kept at 37°C/5% $CO_2$ for 24h in the FourWell plates with fresh media/AA.

The consistency of the 1 kPa hydrogels did not allow SS at 40 dynes/cm$^2$, therefore this condition was discarded. All 14 mechanical conditions were conducted at least in triplicates (technical replicates), making a total of 53 samples (Table S3). Cell alignment on the monolayers was confirmed following SS exposure (Figure S2).

*RNA sequencing*

HUVECs were disrupted using cold QIAzol Lysis Reagent (79306, Qiagen), and RNA was isolated by phenol-chloroform extraction following manufacturer's instructions. RNA concentrations were measured using NanoDrop 2000 (Thermo Scientific). All RNA samples with 260/280 or 260/230 ratios ≤ 1.8 were discarded (average ratios were, respectively, 1.92 and 2.18). RNA sample sequencing was performed by Macrogen Europe (The Netherlands). Samples were subjected to DNAse treatment, a purification step and an RNA quality control using a 2100 Bioanalyzer (Agilent technologies). Libraries were prepared and quality checked for size and quantity by 2100 Bioanalyzer and qPCR, respectively. TruSeq stranded mRNA sequencing was performed using an Illumina NovaSeq 6000 system, with a read length of 100 bp paired end (PE; 60 million reads). Reads were demultiplexed and fastQ files and fastQC[8] reports were obtained from Macrogen. The steps for adapter trimming and alignment were performed in the Linux console. Briefly, sequencing adapters were trimmed from the fastQ files using TrimGalore v.0.6.10 for paired-end samples,[9] and fastQC (version 0.11.9) reports were obtained again. Hisat2 was used to generate map files for the human reference genome (GRCh38 (patch release p14) and to extract splice sites from its GTF annotation file (https://www.ncbi.nlm.nih.gov/assembly/GCF_000001405.40/). The alignment of the reads to the human genome was performed using Hisat2 for paired-end samples[10] and Bam files were obtained. Samtools[11] was used to sort the Bam files and to index these into Bai files. A gene read counts matrix was obtained from these using featureCounts[12] from subread v2.0.3 for paired-end samples. The RNA-Seq dataset can be found in the NCBI Gene Expression Omnibus (GEO) repository with accession number GSE262429.

*Differential expression analysis*

Samples with < 70% assigned reads to genes were discarded (AE, AH, Ai, AX, Y, Z); these contained high levels of ribosomal RNA, which was consistent with their percentage of multi-mapped reads and with FastQC reports (Table S4). On the remaining 47 samples, the percentage of assignment to genes ranged from 89.3% to 70.7%. All ribosomal gene counts were removed from the gene counts matrix (see Table S5 for the excluded ribosomal gene counts).

Exploratory and differential expression analyses on the counts matrix were performed in R using DEseq2.[13] The model was designed as "batch + SS + SFN + SS:SFN", where batch was included to correct for different library preparation and/or sequencing dates. SS and SFN were treated as categorical variables due to the non-linear relationship between SS and SFN. Genes with low counts (< 10) or with less than 2 samples having at least 10 counts were excluded.

Only for visualization purposes, the DESeq2 dds object was normalized using variance-stabilizing transformation (VST),[14] which accounts for size factor estimations (random variability) and for library depth and size. Moreover, for visualization, the batch effect was corrected using limma's function removeBatchEffect.

Principal Component Analysis (PCA) was performed to explore the global variance structure across the 14 conditions. PCA was computed on the VST-corrected counts using the 500 most variable genes, and samples were colored by either SS or SFN groups. Static SS samples formed a distinct cluster separate from all other conditions. To further investigate potential structure among the non-static samples, we applied Uniform Manifold Approximation and Projection (UMAP) on the VST counts after excluding the static SS group. UMAP was run on the top 500 most variable genes (as in PCA) with Euclidean distance metric, number of neighbors = 6, minimal distance = 0.3, and fixed random seed = 42. The resulting two-dimensional coordinates (UMAP1 and UMAP2) were used to visually assess separation among SS and SFN groups. Unsupervised clustering was then applied to the UMAP embeddings. We used k-means clustering on the 2-dimensional UMAP coordinates to identify discrete sample groups within the non-static conditions. The value of $k = 3$ was chosen based on visual inspection of the UMAP structure and biological interpretability.

The 47 samples were then tested for differential expression (DE) analysis (replicates in Table S3). First, DESeq2 was used to perform likelihood ratio tests (LRTs) for SS, substrate SFN, and their interaction. Gene counts were fitted to a full negative binomial generalized linear model containing SS, SFN, and their interaction.  Three reduced models lacking SS, SFN or the interaction term were compared with the full model using LRTs to assess whether including each term improved the ability of the model to explain the observed gene expression counts[15] (see models in Figure 2A). When testing the contribution of a main factor (SS or SFN), the interaction term was also removed from the reduced model to preserve model hierarchy, as interactions should not be retained in the absence of their corresponding main effects. The "batch" factor was kept in all models (full and reduced). *P* values were adjusted using the Benjamini-Hochberg procedure. Genes with adjusted $p < 0.05$ were considered significant, indicating that removal of the tested term significantly worsened model fit. Complete gene lists for each LRT are provided in Supplementary Data 1.

To determine fold changes in gene expression through the various conditions, Wald tests were run in DESeq2, resulting in pair-wise comparisons between each condition and a reference condition. Classically, *in vitro* work has been performed under static conditions and using plastic or glass surfaces; thus, we selected our stiffest static condition as the reference (100 kPa at static). We extracted fold changes and adj-*p* for all genes and built up a DE data set containing the LRT results for each term and the fold changes and Wald test adj-*p* results across all the condition contrasts for all genes. Complete gene lists for each Wald test result can be found in Supplementary Data 2-3.

#### *Gene expression functional analyses - IPA*

Using CORE analyses by Ingenuity Pathway Analysis (IPA; version 107193442, Genomics Core KU Leuven), we explored the dynamics in biological processes behind the gene lists and their fold changes. *Cardiovascular System* showed as the top *Physiological System Development and Function* for the LRT interaction list. From this, the top significant biological processes were selected and displayed as networks. Then, fold change data (from contrasts) was overlaid on the displayed network to visualize the processes

Z-scores predicting their activation or inactivation, as reported by the curated database in IPA. Briefly, Z-scores are correlations between gene fold changes and literature-based effects of those genes on specific processes or functions (inhibiting or enhancing). Positive Z-scores would predict activation, and correspond to positive fold changes from enhancing genes, or negative fold changes from inhibiting genes; negative Z-scores would predict inactivation, and correspond to positive fold changes from inhibiting genes, or negative fold changes from enhancing genes.

Another DE dataset was obtained containing the LRT results for each term and the fold changes and Wald test adj-*p* results for contrasts against the own static conditions. IPA CORE analyses and comparisons were run as described.

To investigate whether IPA-identified process activation patterns were reflected in the UMAP structure, we overlaid angiogenesis activation Z-scores onto the UMAP embedding of non-static samples. Because IPA provides one Z-score per experimental condition, whereas UMAP represents individual samples, all samples were assigned the Z-score corresponding to their condition. The IPA activation color gradient was applied to the UMAP embedding, enabling visualization of whether angiogenesis activation increased or decreased across specific UMAP regions. To summarize trends at the condition level, UMAP centroids were computed for each condition. To test whether UMAP-derived clusters captured variation in angiogenesis activation, k-means clusters assignments were used to compare IPA Z-scores across clusters. Differences in Z-score distributions were visualized using violin plots with overlaid boxplots, and statistical significance was assessed using one-way ANOVA with IPA Z-score as the response and cluster as the factor.

*Gene expression functional analyses - KEGG*

Because KEGG relies on a distinct annotation resource and analytical framework compared with IPA, KEGG-based pathway analysis was performed as an independent validation of the IPA results. KEGG gene-to-pathway relationships were retrieved using KEGGREST. For each comparison, pathway-level scores were calculated as the mean $log_2$FC of genes belonging to each KEGG pathway. To focus on biologically responsive pathway members, only genes significantly differentially expressed in at least one of the two conditions being compared were retained (union criterion). Pathways represented by fewer than three contributing genes were excluded. KEGG pathways were further restricted to the top-level categories “Cellular Processes” and “Environmental Information Processing”. For visualization, pathways related to “caffeine”, “gentamicin”, “xenobiotic”, “drug metabolism”, “carcinogenesis” and “antibiotic” were excluded.

Pathway-level relationships were assessed through pairwise comparisons of KEGG pathway scores between 10 kPa and 100 kPa under each SS condition: 25 dynes/cm$^2$ (r = 0.87, $p$ = 4.8x10$^{-18}$) and at 40 dynes/cm$^2$ (r = 0.71, $p$ = 9.6x10$^{-10}$). To visualize differential pathway regulation, KEGG scores from each pairwise comparison were plotted in scatter plots, where an identity line (y = x) indicates equal pathway activation; pathways above or below this line correspond to processes preferentially activated at 100 kPa or 10 kPa SFN, respectively. To quantify SFN-dependent pathway divergences within each SS condition, ΔSFN scores were calculated for each KEGG pathway as ΔSFN = mean $\log_2$FC(100 kPa) – mean $\log_2$FC(10 kPa). These values were visualized using ordered bar plots for each SS condition, accompanied by density distributions to illustrate the global directional bias of pathway activation relative to zero; positive or negative ΔSFN values correspond to pathways with relatively higher activation at 100 kPa or 10 kPa, respectively.

During KEGG pathway analysis, "Neuroactive ligand–receptor interaction" (hsa04080) and "Neuroactive ligand signaling" (hsa04082) pathways were identified among the top-ranked pathways. Gene overlap between these pathways was partial (Jaccard index ≈ 0.30), and their pathway-level responses differed across conditions, supporting their treatment as distinct modules. KEGG pathway-level analyses (including scatter and bar plots) were performed using the full gene sets defined by KEGG. For clarity in visualization, these pathways were relabeled as "Ligand–receptor signaling (A; hsa04080)" and "Ligand–receptor signaling (B; hsa04082)". To aid biological interpretation, gene composition within hsa04080 was further inspected, revealing a subset of ligand–receptor signaling genes with established vascular relevance. The vascular subset reproduced the direction of change observed for the full hsa04080 pathway across conditions, indicating that pathway-level behavior was partially driven by vascular-relevant components. This subset was not used for pathway-level scoring but was retained for downstream gene-level analyses and interpretation.

To identify gene-level drivers of the divergence between the two angiogenic-activated states (25 dynes/cm$^2$ at 100 kPa vs. 40 dynes/cm$^2$ at 10 kPa), a derived contrast was calculated for each gene as Δmech = $\log_2$FC (40 dynes/cm$^2$, 10 kPa) − $\log_2$FC (25 dynes/cm$^2$, 100 kPa). Positive or negative Δmech values indicate genes with relatively higher expression in the 40 dynes/cm$^2$ at 10 kPa condition or the 25 dynes/cm$^2$ at 100 kPa condition, respectively. Genes were ranked by Δmech values, and the candidate universe was restricted to genes belonging to KEGG pathway families identified as relevant in the preceding pathway-level analysis. For visualization, three functional groups were defined by combining pathway-derived candidate sets: specifically, "adaptive/remodeling response" combined genes driving AMPK, HIF, FoxO, ECM, and the vascular submodule derived from the Neuroactive ligand-receptor interaction pathway (hsa04080); "inflammatory/immune-adhesive response" combined genes driving NFκB, TNF, cytokine-cytokine, CAM, and JAK/STAT pathways; and "stress/damage response" combined genes driving ferroptosis, apoptosis, p53 and cellular senescence pathways. Genes were assigned to these categories based on pathway membership, and poorly annotated loci, antisense transcripts, and pseudogenes were excluded for interpretability. From this candidate universe, top positive and negative Δmech genes were inspected, and a non-redundant subset of genes with strong effect sizes and clear biological relevance was manually curated for visualization. KEGG pathways ranked by ΔSFN and gene contributing to the divergence between angiogenic profiles ranked by ΔMech are provided in Supplementary Data 4.

*In vitro EC migration experiments*

Hydrogels were prepared similarly as previously described. In this case, cover glasses were treated with APTS and glutaraldehyde. Hydrogel substrates of 0.5, 3.2, 4.5, 10 and 35 kPa were prepared by mixing 40% acrylamide and 2% bis-acrylamide to final concentrations depicted in Table S6. Afterwards, 10 µL of 10% APS and 0.8 µL of TEMED were added to 1 mL Bis/Acrylamide. The mixture was then added onto the cover glass, covered with a Sigmacote®-treated coverslip, and let rest for 30 min at RT. Hydrogels were functionalized as previously described, but were coated with 15 µg/mL fibronectin (Corning, 354008) for 2h at 37°C. Passage 5 HUVECs, 15% of which were dyed using CellTrackerTM Green CMFDA Dye (ThermoFisher, C2925), were seeded on fibronectin-coated hydrogels at a density of 1,600 cells/mm$^2$. Cells were cultured for 30h in EGM-2MV media at 37°C to allow them to attach and become fully confluent. Afterwards, cells were starved for 11h (EGM-2MV with only 0.5% FBS instead of 5% and without VEGF and FGF), and subjected to SS levels of 1 dyne/cm$^2$ for 5h while being imaged once every 30 min on Zeiss LSM 700 AxioObserver confocal microscope. After acquisition, spots creation and tracking algorithm of Imaris 9.1 (Bitplane) software was used to track the dyed cells (as previously described).[16] Migration rates (µm/h) were calculated for each traced cell by summing up the covered distances registered at each timepoint and dividing them by the time lapse of each experiment.

*Active YAP1 staining*

HUVECs were cultured on functionalized hydrogels at 1, 10, and 100 kPa, as detailed in previous sections. After 24h of SS at 0, 5, 15, 25, and 40 dynes/cm$^2$, slides were fixed in 4% PFA for 15 min at 4°C, washed and preserved in PBS. Gels were cut into approximately 0.5 cm$^2$ pieces and placed in 24-well plates. All steps during the staining protocol were performed under gentle agitation. Gels were equilibrated in Tris buffer (50 mM Tris, 137 mM NaCl, 2.7 mM KCl, pH 7.5) containing 0.05 % Triton x100 (TBST) during 3 washes of 15 min at RT. Gels were blocked and permeabilized for 1h in blocking solution (Tris buffer + 0.5% blocking reagent TSA-kit, Perkin Elmer) containing 0.2% Triton x100 (TNBT). Gels were stained overnight at 4°C in a

1/500 dilution in TNBT of anti-active YAP1 antibody [EPR19812] (AB205270, Abcam). Gels were washed 3 times in TBST for 15 min and incubated for 2h at RT in a 1/250 dilution of TO-PRO™-3 Iodide (642/661) (T3605, Thermo Fisher Scientific) and a 1/500 dilution of anti-rabbit-AF488 antibody (A11008, Thermo Fisher Scientific) in TBST. Gels were washed 3 times in TBST for 15 min and were mounted on slides using ProLong Gold antifade mountant (P36930, Invitrogen). Coverslips were sealed with nail polish and slides were kept in the dark at 4°C overnight, after which cells were imaged. Three hydrogel pieces were imaged per condition, when possible; otherwise, images were taken from different areas of a hydrogel piece.

YAP1 intensity was quantified using ZEN 3.9 Lite software (ZEISS). Imaging exposure settings were kept constant across all imaging sessions. Regions of interest (ROIs) were defined for nuclei (TO-PRO channel) and cell perimeter (YAP1 channel). Mean intensities were extracted, and relative nuclear YAP1 intensity was calculated as the ratio of nuclear to the whole cell value (N/C). A minimum of 23 nuclei were measured per condition (39 nuclei per condition on average). Mean YAP1 nuclear intensity was calculated for each SS-SFN combination and visualized as a heatmap to highlight global trends across mechanical conditions.

#### *Correlation of interaction-responsive gene expression with YAP1 nuclear localization*

To identify interaction-regulated genes associated with YAP1 nuclear localization, genes significant in the RNA-seq LRT interaction analysis were retained as the starting set, and their gene expression was tested for correlation with YAP1 nuclear localization. YAP1 nuclear localization differed substantially between substrate SFN groups, with baseline relative nuclear intensity values on 1 kPa (~0.69) being markedly lower than those observed on 10 kPa (~1.99) and 100 kPa (~1.70). To prevent these SFN-dependent baseline differences from dominating the correlation analysis, both gene expression (DESeq2 contrasts) and YAP1 localization were expressed relative to the matched static condition within each SFN group. Thus, static conditions therefore had a normalized value of 1 and were excluded from the correlation analysis. Pearson correlations were then calculated between normalized gene expression changes and normalized YAP1 nuclear localization across the 11 non-static conditions. This approach enabled assessment of how SS-induced changes in gene expression covaried with SS-induced changes in YAP1 localization within distinct SFN contexts. Genes with significant Pearson correlations ($|r| > 0.6$, $p < 0.05$) were considered YAP-associated candidate genes. From this set, five genes (*CLEC3B*, *GAB1*, *MATN3*, *MGLL*, and *KCTD20*) were selected for exploratory follow-up to represent both positively and negatively associated candidates. Correlation values for interaction genes and selected candidate genes provided in Supplementary Data 5.

#### *YAP1 knockdown in HUVECs and RT-qPCR*

HUVECs were electroporated with ON-TARGETplus Human *YAP1* siRNA Smartpool (L-012200-00-0005, Dharmacon; *siYap1*) or ON-TARGETplus Non-targeting Pool (D-001810-10-05, Dharmacon; NT). Briefly, 3.5 million cells were resuspended in 100 µL DPBS. A 10 µM dilution of *siYAP1* or NT in 1x siRNA buffer (B-002000-UB-100, Horizon Discovery) was added and cells were transferred to Nucleocuvette™ Vessels from the P5 Primary Cell 4D-Nucleofector™ X Kit (V4XP-5024, Lonza) and electroporated in a 4D-Nucleofector station (Lonza) using the P5 CA-167 program. Endothelial media/AA was immediately added and cells were seeded onto 1 or 100 kPa hydrogels functionalized with collagen type I, at a density of 0.6 million cells per hydrogel. After an overnight incubation at 37°C/5 % $CO_2$, confluent monolayers were exposed to static or 15 dynes/cm$^2$ SS levels for 24 h, after which RNA was extracted.

Real Time quantitative PCR (RT-qPCR) was run for *GAPDH* (as housekeeping gene), *YAP1*, *MGLL*, *MATN3*, *CLEC3B*, *KCTD20*, and *GAB1* using in-house-designed primers (Integrated DNA Technologies; Table S7). Technical replicate Ct values were converted to ΔCt values by subtracting GAPDH Ct from target-gene Ct, and were then averaged to obtain one ΔCt value per biological replicate for each gene and condition; all statistical analyses were performed on these biological replicate ΔCt values. *YAP1* knockdown effects (ΔΔCt) were quantified within each mechanical condition as the difference in mean ΔCt between *siYAP1* and NT groups (*siYAP1* − NT), with 95% confidence intervals estimated by bootstrap resampling. Positive values indicate decreased expression after *YAP1* knockdown, whereas negative values indicate increased expression. See Supplementary Data 5 for replicate numbers, ΔCt, ΔΔCt, and confidence interval values.

#### *Statistical Analysis*

Statistical analyses were all run in R (v. R-4.3.1). The RNA-Seq analysis was conducted on 47 samples ($n$ = 3 to 5 samples per condition; $n$ = 2 for 1 kPa at 15 dynes/cm$^2$) using the LRT and Wald test methods in DESeq2 as previously described. P-values were adjusted for multiple testing using the Benjamini-Hochberg method, and adjusted $p$-values < 0.05 were considered statistically significant. Quadratic and linear regression curves on gene count plots were obtained using the geom_smooth function in ggplot2 in R.

Gene expression functional analyses were run in IPA (v. 107193442) and KEGG. Prediction statistical details for IPA are available online. KEGG pathway analyses were based on pathway-level aggregation of $\log_2$ fold changes from significant pathway-member genes, using adjusted $p < 0.05$ and a minimum pathway size of 3 genes.

Differences in YAP1 localization were calculated using two-way ANOVA followed by Tukey's honestly significant difference (HSD) post hoc test. For qPCR experiments, statistical analyses were performed on ΔCt averaged at the biological replicate level. *YAP1* knockdown effects were quantified as differences between *siYAP1* and NT control within each condition. Because the experiment was exploratory and sample sizes were small (*n* = 3 – 4 biological replicates per condition), effect sizes were summarized using bootstrap resampling to estimate 95% confidence intervals.

### Supplementary Materials References


1. Tse JR, Engler AJ. Preparation of hydrogel substrates with tunable mechanical properties. *Curr Protoc Cell Biol* 2010:1–16.
2. Bastounis EE, Ortega FE, Serrano R, Theriot JA. A multi-well format polyacrylamide-based assay for studying the effect of extracellular matrix stiffness on the bacterial infection of adherent cells. *Journal of Visualized Experiments* 2018;**2018**.
3. Judokusumo E, Tabdanov E, Kumari S, Dustin ML, Kam LC. Mechanosensing in T lymphocyte activation. *Biophys J* 2012;**102**:L5–L7.
4. Shrestha KR, Lee DH, Chung W, Lee S-W, Lee Y, Yoo SY. Biomimetic virus-based soft niche for ischemic diseases. *Biomaterials* 2022;**288**:121747.
5. Huynh J, Nishimura N, Rana K, Peloquin JM, Califano JP, Montague CR, et al. Age-Related Intimal Stiffening Enhances Endothelial Permeability and Leukocyte Transmigration. *Sci Transl Med* 2011;**3**:112ra122.
6. Ng MR, Besser A, Danuser G, Brugge JS. Substrate stiffness regulates cadherin-dependent collective migration through myosin-II contractility. *J Cell Biol* 2012;**199**:545.
7. Rouillard AD, Gundersen GW, Fernandez NF, Wang Z, Monteiro CD, McDermott MG, et al. The harmonizome: a collection of processed datasets gathered to serve and mine knowledge about genes and proteins. *Database (Oxford)* 2016;**pii**.
8. Andrews S. FastQC: A Quality Control Tool for High Throughput Sequence Data [Online]
9. Krueger F, James F, Ewels P, Afyounian E, Weinstein M, Schuster-Boeckler B, et al. FelixKrueger/TrimGalore: v0.6.10 - add default decompression path
10. Kim D, Paggi JM, Park C, Bennett C, Salzberg SL. Graph-based genome alignment and genotyping with HISAT2 and HISAT-genotype. *Nature Biotechnology 2019 37:8* 2019;**37**:907–915.
11. Danecek P, Bonfield JK, Liddle J, Marshall J, Ohan V, Pollard MO, et al. Twelve years of SAMtools and BCFtools. *Gigascience* 2021;**10**:1–4.
12. Liao Y, Smyth GK, Shi W. featureCounts: an efficient general purpose program for assigning sequence reads to genomic features. *Bioinformatics* 2014;**30**:923–930.
13. Love MI, Huber W, Anders S. Moderated estimation of fold change and dispersion for RNA-seq data with DESeq2. *Genome Biol* 2014;**15**:1–21.
14. Anders S, Huber W. Differential expression analysis for sequence count data. *Genome Biol* 2010;**11**:1–12.
15. Love MI, Anders S, Huber W. Analyzing RNA-seq data with DESeq2 - Likelihood Ratio Test. *https://bioconductor.org/packages/devel/bioc/vignettes/DESeq2/inst/doc/DESeq2.html#likelihood-ratio-test*.
16. Tabibian A, Ghaffari S, Vargas DA, Oosterwyck H Van, Jones EAV. Simulating flow induced migration in vascular remodelling. *PLoS Comput Biol* 2020;**16**.
